# Elevated cholesterol in APOE4 astrocytes drives mitochondrial cristae collapse and ATP synthase dysfunction

**DOI:** 10.64898/2026.08.04.742391

**Authors:** Seungtae Lee, Mikel Munoz-Oreja, Marina Villar-Fernandez, Leire Goicoechea-Barrenechea, Uxoa Fernandez-Pelayo, Diego Perez-Rodriguez, Matthew Gegg, Amaia Lopez de Arbina, Antonella Spinazzola, Ian James Holt

**Affiliations:** Biogipuzkoa Health Research Institute, 20014, San Sebastián, Spain; CIBERNED (Center for Networked Biomedical Research on Neurodegenerative Diseases, Ministry of Economy and Competitiveness, Institute Carlos III, 28031 Madrid, Spain; Department of Clinical and Movement Neurosciences, UCL Queen Square Institute of Neurology, Royal Free Campus, London NW3 2PF, UK; IKERBASQUE, Basque Foundation for Science, 48013 Bilbao, Spain; Universidad de País Vasco, Barrio Sarriena s/n, 48940, Leioa, Bilbao, Spain

**Keywords:** mitochondria, cholesterol, ATP synthase, cristae, astrocytes, APOE, cyclodextrin

## Abstract

Cholesterol imbalance is a hallmark of major human diseases, including atherosclerosis and Alzheimer’s disease, both of which are also associated with mitochondrial dysfunction, yet the mechanistic links between cholesterol and mitochondria remain poorly understood. Here we show that elevated intracellular cholesterol in murine astrocytes expressing the Alzheimer’s disease risk variant *APOE4* disrupts the inner mitochondrial membrane, manifesting as sparse, truncated cristae alongside an excess of cristae junction complexes. These structural abnormalities are accompanied by loss of respiratory chain complexes I and IV, and reduced respiration; nevertheless, low proton leak and reverse ATP synthase activity combine to generate an elevated mitochondrial membrane potential. Strikingly, *APOE4* astrocytes are hypersensitive to the ATP synthase inhibitor oligomycin, demonstrating a profound dysfunction of the enzyme, and cholesterol sequestration with methyl-β-cyclodextrin abrogates this toxicity, establishing elevated cholesterol as its proximate cause. Another method of targeting ATP synthase, epicatechin, which prevents the enzyme operating in reverse, attenuated both cholesterol- and respiratory chain inhibitor-induced cell death. Reciprocally, reducing intracellular cholesterol via nutrient restriction restored cristae architecture and partially rescued respiratory chain complex abundance. These findings identify mitochondrial cholesterol as a critical determinant of cristae architecture and ATP synthase function and suggest that cholesterol-driven mitochondrial dysfunction may be a unifying feature of cholesterol-related disorders from neurodegeneration to atherosclerosis.

**GRAPHICAL ABSTRACT:** Elevated intracellular cholesterol in *APOE4* astrocytes disrupts the inner mitochondrial membrane and ATP synthase, resulting in sparse cristae, increased ATP hydrolysis and heightened oligomycin sensitivity. The vulnerability to oligomycin is abrogated by cholesterol sequestration with cyclodextrin, indicating causality. These findings position cholesterol as a key determinant of the cristae landscape and ATP synthase function, highlighting the importance of the emerging field of mitochondrial-cholesterol crosstalk.

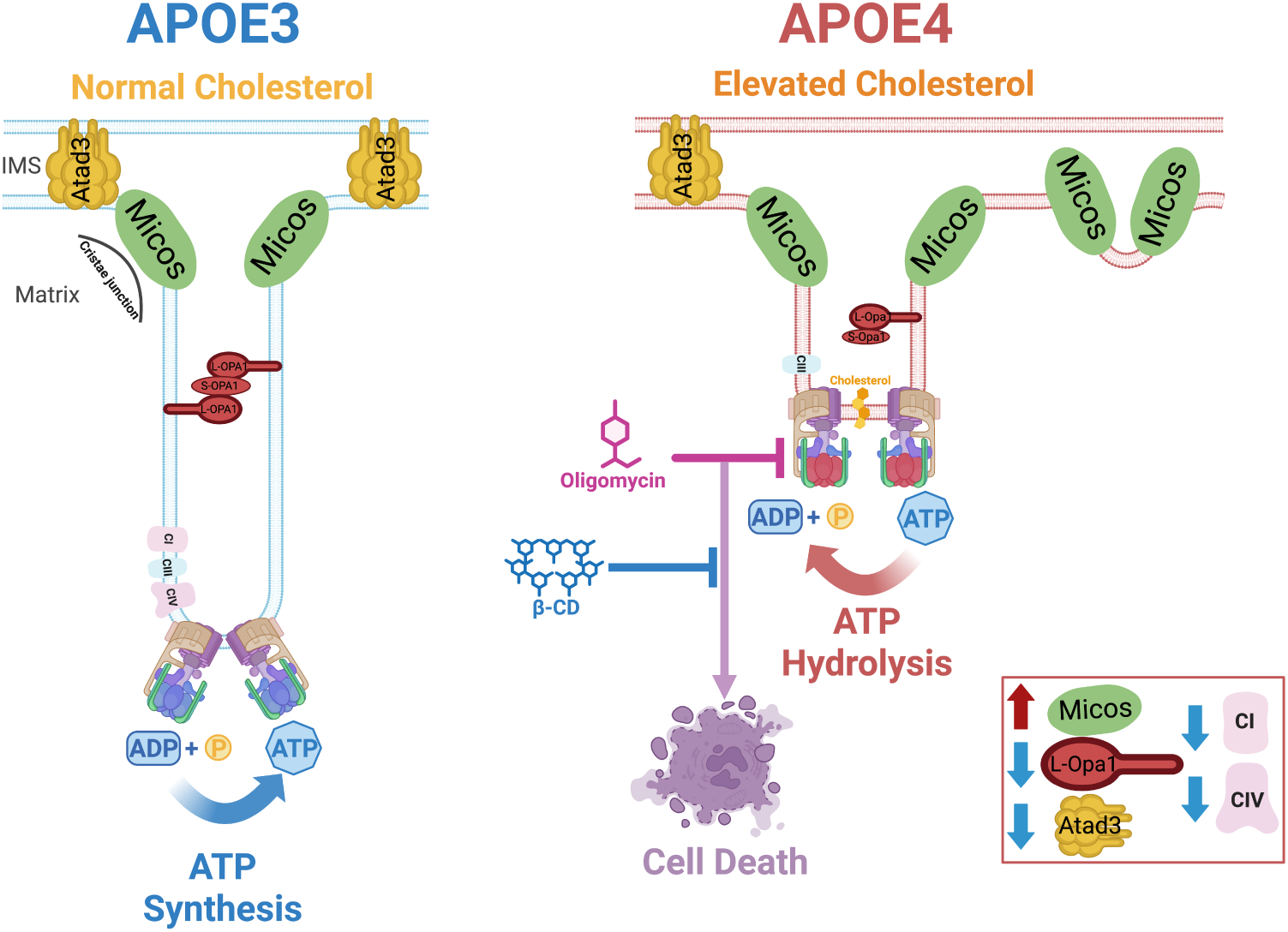

## INTRODUCTION

Cholesterol is a critical structural component of most cell membranes. It is organized in platforms or microdomains (aka lipid rafts) that influence numerous membrane activities (Doole et al., 2022; Lingwood & Simons, 2010; Sezgin et al., 2017). Perturbed cholesterol homeostasis is of major importance to human health through its links to atherosclerosis and Alzheimeŕs, and other neurodegenerative, diseases (Duan et al., 2022; Fortea et al., 2024). Alzheimeŕs disease is also associated with mitochondrial dysfunction at an early stage (Yao et al., 2009; Venkataraman et al., 2022), yet whether defective cholesterol and mitochondrial metabolism share a common etiology has barely been explored. This possibility has increasing resonance as mitochondria have emerged in recent years as important players in cholesterol homeostasis, via studies of the mitochondrial trans-membrane protein ATAD3 (ATPase family AAA domain-containing protein 3). Fibroblasts from patients carrying pathological *ATAD3* variants and an *Atad3* conditional knockout mouse display reprogrammed cholesterol metabolism (Peralta et al., 2018); Desai et al., 2017; Gunning et al., 2020; Kiesel et al., 2025; Muñoz-Oreja et al., 2024). The cholesterol-related abnormalities of *ATAD3* mutant cells overlap those of the cholesterol trafficking disorder Niemann Pick type C disease (NPC) (Desai et al., 2017; Muñoz-Oreja et al., 2024); and reciprocally, *NPC* mutant cells and knockout mice display mitochondrial abnormalities (Goicoechea et al., 2024; Höglinger et al., 2019; Pfrieger, 2023). These findings point to a firm mechanistic link between cellular cholesterol metabolism and mitochondria and suggest that cholesterol imbalance impairs mitochondrial function. If these findings generalize, then essentially all forms of cholesterol imbalance should provoke mitochondrial dysfunction.

Apolipoprotein E (APOE) plays a central role in cellular cholesterol trafficking, and the *APOE4* variant is associated with elevated intracellular cholesterol and greatly elevated risk of Alzheimer’s disease (de Leeuw et al., 2022; TCW et al., 2022). Hence, *APOE* models can serve as test beds for investigating cholesterol-mitochondrial crosstalk. Nowhere is APOE more important than in astrocytes, as they are the major producers of brain cholesterol, which they package into APOE-containing lipoproteins for transport to neurons (Mahley, 2016); and there are already indications that *APOE4* astrocytes exhibit altered mitochondrial dynamics and impaired oxidative phosphorylation (Lee et al., 2023; Schmukler et al., 2020). However, the structural and mechanistic bases of these changes were unclear; nor was it clear to what extent cholesterol was responsible for the mitochondrial phenotypes. Therefore, we investigated the mechanistic link between cholesterol metabolism and mitochondrial function in murine astrocytes carrying human *APOE3* and *APOE4* variants that display altered cholesterol homeostasis (Staurenghi et al., 2022). Our analysis reveals elevated intracellular cholesterol disrupts the inner mitochondrial membrane and ATP synthase - abnormalities that may explain the early appearance of mitochondrial dysfunction in cholesterol-related diseases.

## RESULTS

### *APOE4* rewires cholesterol handling

After confirming variant-specific *APOE* expression by Western blot (**Figure 1A**), we began to assess how *APOE* variants affect mitochondrial-cholesterol crosstalk by evaluating cholesterol homeostasis. The plasma membrane cholesterol transporter ABCA1 is a key regulator of intracellular cholesterol that is depressed in human *APOE4* astrocytes (TCW et al., 2022). Likewise, murine astrocytes expressing human *APOE4* had a fifth of the AbcA1 of *APOE3* astrocytes (**Figure 1B, 1C**); together with 4.5-fold more Hmgcr cleavage product (**Figure 1D, 1E**), another indicator of excess intracellular cholesterol (Tsai et al., 2012). We next measured Gm1 ganglioside levels, as GM1 promotes tight cholesterol packing and reduced membrane fluidity (Galimzyanov et al., 2017), and it binds preferentially to APOE (Zhang et al., 2025). Gm1 levels were six-fold higher in *APOE4* than *APOE3* astrocytes (**Figure 1F, 1G**). Thus, expression of human *APOE4* in murine astrocytes markedly perturbs cholesterol homeostasis. Given the altered intracellular cholesterol state, we tested sensitivity to exogenous cholesterol. APOE4 astrocytes were approximately three times more sensitive to cholesterol toxicity than APOE3 astrocytes, with 10 µg/mL cholesterol reducing cell numbers by ~80% over 48 hours (**Figure 1H, 1I**). The starkest differences appeared between 24 and 48 hours where the number of APOE3 astrocytes increased almost three-fold, whereas it halved for *APOE4* astrocytes, with the latter result indicating cell death, not merely growth arrest (**Figure S1**). Together these findings indicate that *APOE4* remodels cholesterol homeostasis in murine astrocytes rendering them highly vulnerable to exogenous cholesterol.

**Figure 1.**
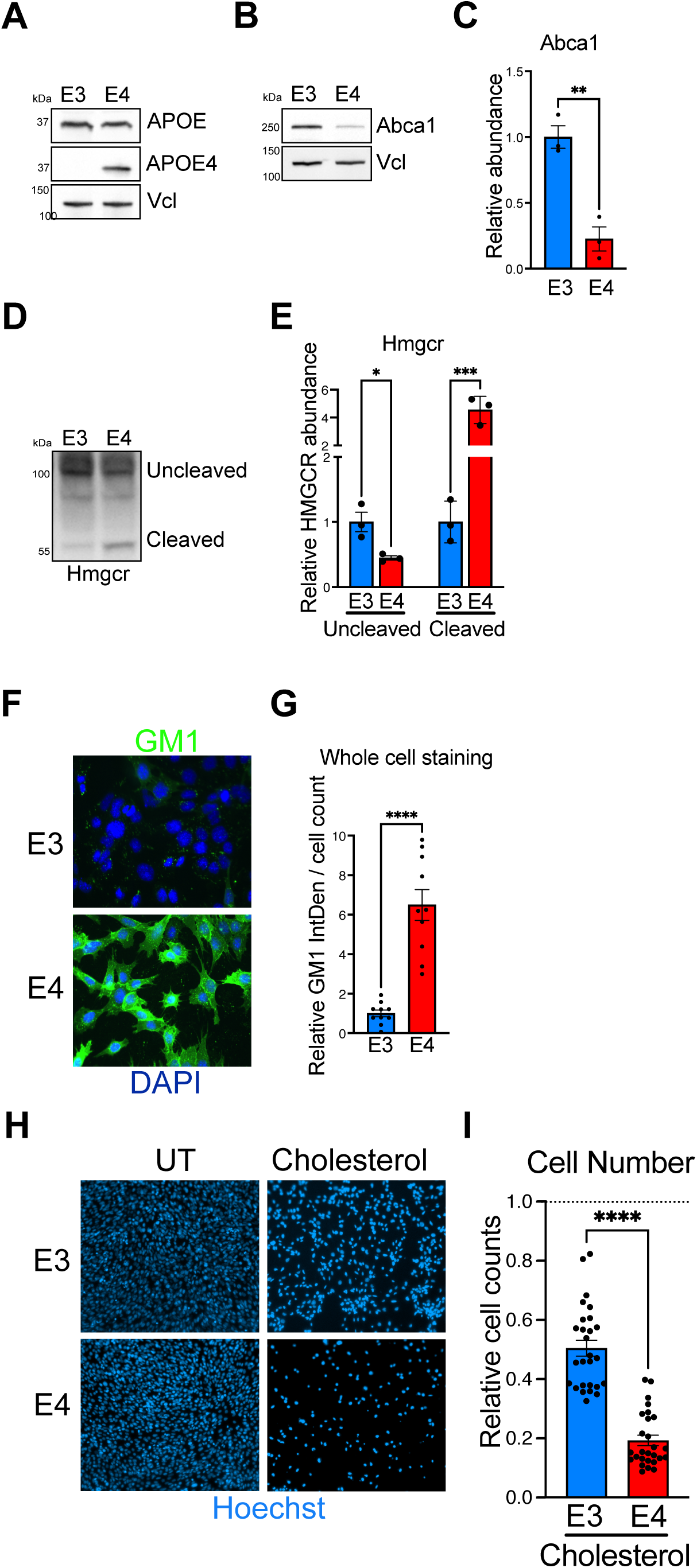
*APOE4* astrocytes have elevated intracellular cholesterol, rendering them intolerant of cholesterol supplementation. Isolated protein from humanized *APOE3* (E3) and *APOE4* (E4) astrocytes were fractionated by SDS-PAGE and after transfer to solid support immunoprobed for **A)** APOE and APOE4; **B)** the plasma membrane cholesterol transporter, Abca1, quantified in C, n = 3, p = 0.003 and **D)** the rate-limiting enzyme of cholesterol biosynthesis Hmgcr, all with Vinculin as a loading control, with quantification in **E** (n = 3 independent experiments, p = 0.02 and p = 0.0002 for uncleaved and cleaved Hmgcr, respectively). **F**) *APOE3* and *APOE4* astrocytes immunostained for Gm1 ganglioside (green) and DAPI stained DNA (blue). **G**) Gm1 quantification from n = 5 independent experiments, p = 0.0001. **H**) astrocytes treated without (UT) and with 10 µg/mL cholesterol for 48 hours, and the nuclei stained with Hoechst. **I**) Quantification of Hoechst-stained cells from 8 independent experiments, with the number of cholesterol-treated astrocytes normalized to the number of untreated cells grown in parallel in otherwise identical conditions, p = 0.0001. NB Plating fewer cells in smaller areas, as occurred in several subsequent experiments, accelerated or accentuated the toxic effects of cholesterol supplementation and some other treatments, and so cannot be directly compared.

### The mitochondria of APOE4 astrocytes have fewer and shorter cristae but more cristae junction complexes than those of APOE3 astrocytes

We expected the elevated intracellular cholesterol of the APOE4 astrocytes to adversely impact membrane composition and architecture, and that the highly folded inner mitochondrial membrane (cristae) would be among the most affected. Ultrastructural analysis via electron microscopy revealed that *APOE4* mitochondria have fewer cristae per organelle, including over a quarter lacking cristae entirely, whereas all *APOE3* mitochondria contained cristae; and the average cristae length was twice that of *APOE4* mitochondria (**Figure 2A-2D**). Concordant with the electron micrographs, the long isoforms of Opa1 that support cristae (Del Dotto et al., 2017; Hu et al., 2020) were 3.5 times less abundant in *APOE4* than *APOE3* astrocytes (**Figure 2E, 2F**).

**Figure 2.**
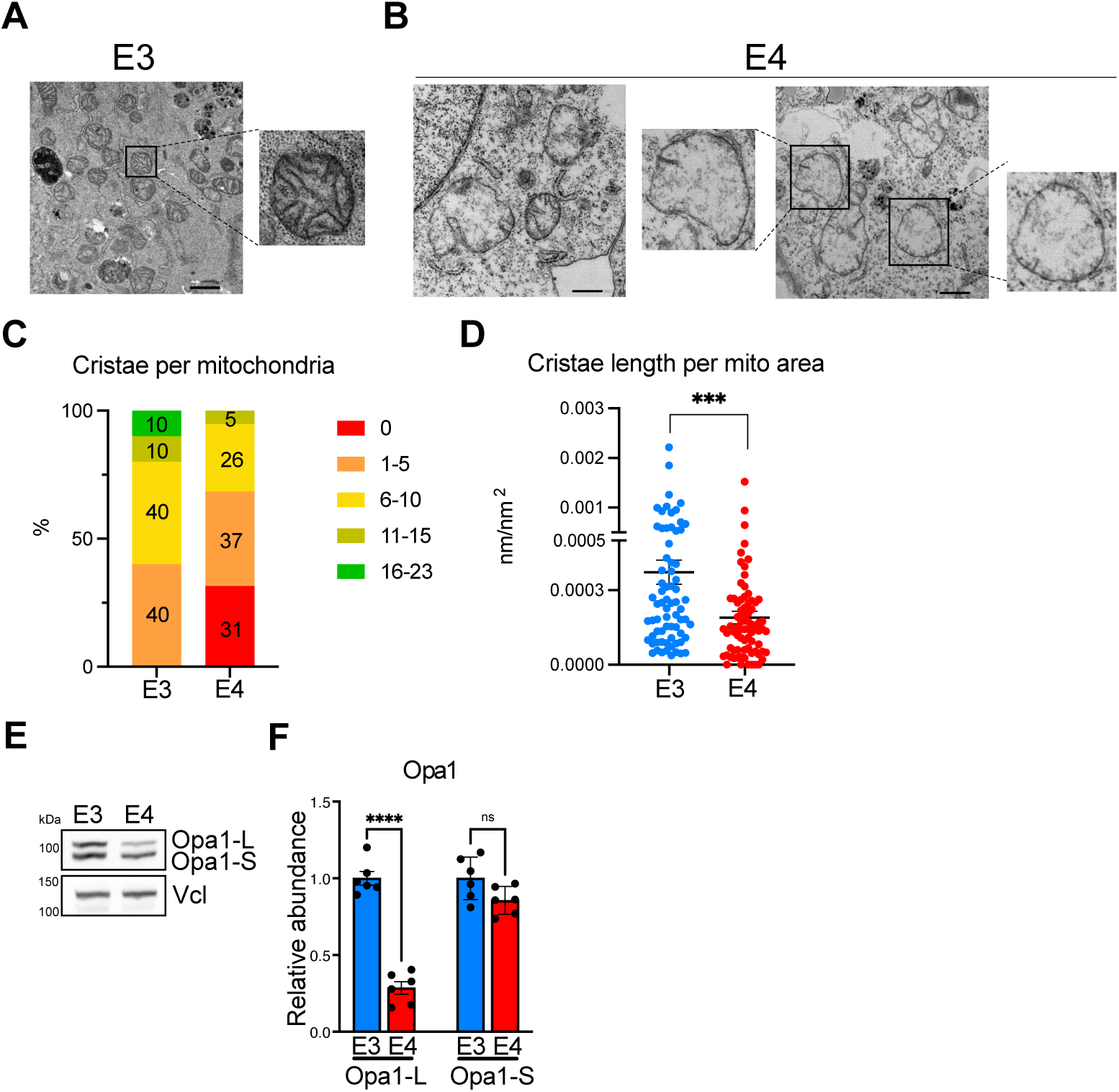
*APOE4* astrocytes have abnormal mitochondrial ultrastructure with sparse, truncated cristae. Electron micrographs of murine astrocytes carrying the **A**) *APOE3* (E3), or **B**) *APOE4* (E4) variant, with zoomed images highlighting individual mitochondria. Scale bars from left to right = 1 μm, 500 nm and 1 μm. Panels **C** and **D**, charts indicating the differences between mitochondrial cristae number and length between *APOE3* and *APOE4* astrocytes. **E**) Opa1 long (Opa1-L) and short (Opa1-S) isoform abundance in *APOE3* and *APOE4* astrocytes via immunoblotting. **F**) Quantification of the long form of Opa1; there was a 3.5-fold difference in abundance between E4 and E3 astrocytes (p = 0.0001, n = 6 experiments).

Because cristae junction complexes are critical to cristae maintenance (Del Dotto et al., 2017; Hu et al., 2020), we expected these complexes to be scarce in the *APOE4* astrocytes. On the contrary, the abundance of the core cristae junction component Mic60/mitofilin, detected as a denatured protein or in the fully assembled Micos and Mib complexes of 720 and 1400 kDa, respectively, was 2.2-fold higher in *APOE4* than *APOE3* astrocytes (**Figure 3A-3C**). Thus, we infer that impaired cristae formation is downstream of cristae junction complex assembly, or that excess cholesterol in the mitochondria causes cristae collapse.

**Figure 3.**
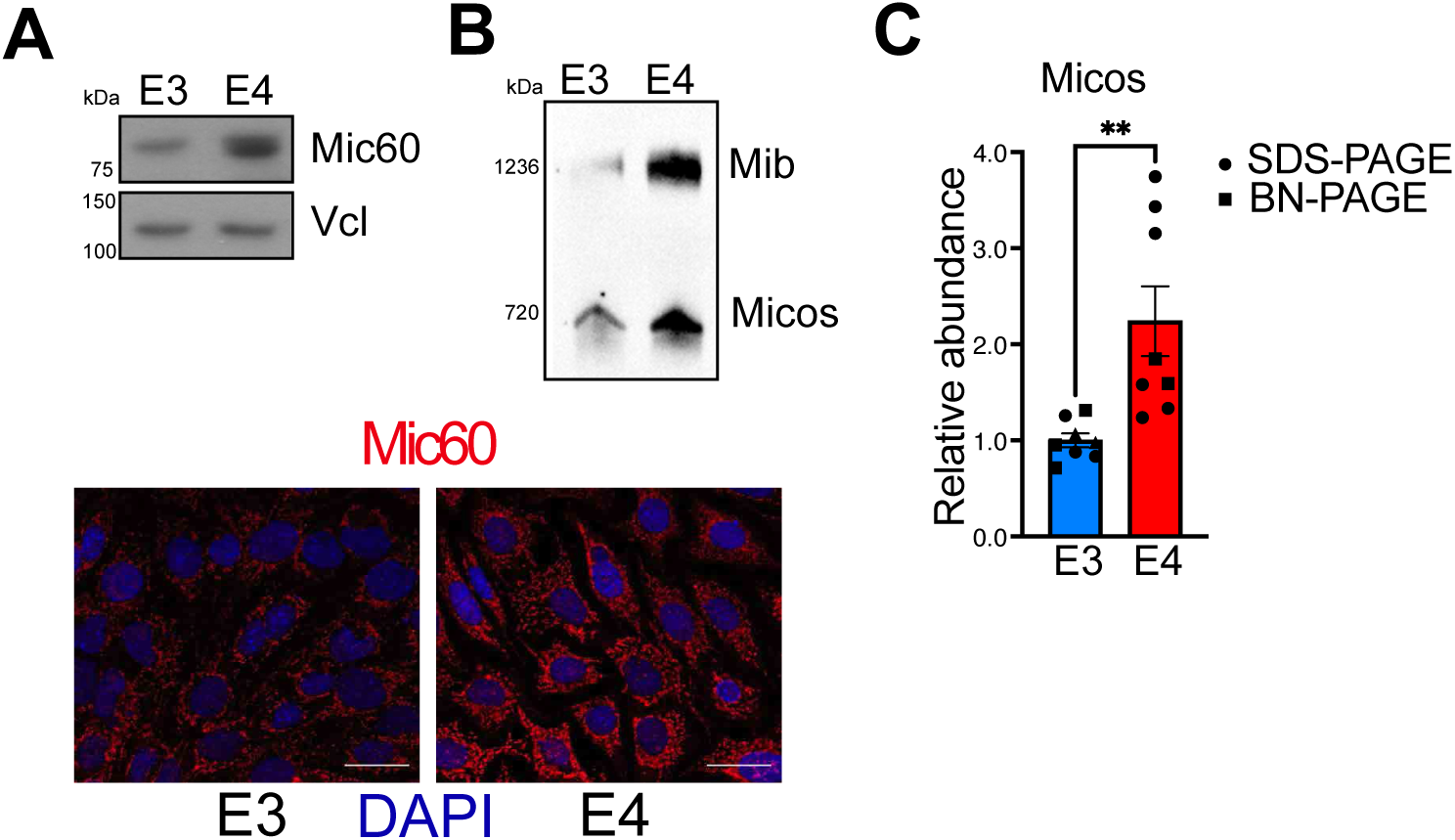
*APOE4* astrocytes have a surfeit of cristae junctions. **A**) Mic60/Mitofilin in *APOE3* and *APOE4* astrocytes detected by immunocytochemistry or after fractionation of cellular protein via SDS-PAGE, or **B**) native gel electrophoresis (BNE), both quantified in **C**); n = 8 experiments, p = 0.005.

### APOE4 is associated with extensive remodeling of the Oxidative Phosphorylation System

The highly folded inner mitochondrial membrane is designed to accommodate the millions of respiratory chain and ATP synthase complexes required to meet the cell’s energy demands. Hence, the disrupted cristae were expected to impact oxidative phosphorylation. Concordantly, the APOE4 astrocytes had lower levels of respiratory chain complexes I and IV (**Figure 4A, 4B**); and lower respiration than their *APOE3* counterparts (**Figure 4C, 4D**), like *APOE4* human astrocytes (Lee et al., 2023). On the other hand, the abundance of complex III was unaffected, and ATP synthase was more abundant in *APOE4* than *APOE3* astrocytes (**Figure 4A, 4B**). The altered membranes and oxidative phosphorylation system increased the vulnerability of *APOE4* astrocytes to mitochondrial inhibitors. Forty-eight hours exposure to 25 nM antimycin A, a respiratory complex III inhibitor, or 5 nM oligomycin, that inhibits ATP synthase, were both twice as toxic to *APOE4* than to *APOE3* astrocytes (**Figure 4E, 4F**).

**Figure 4.**
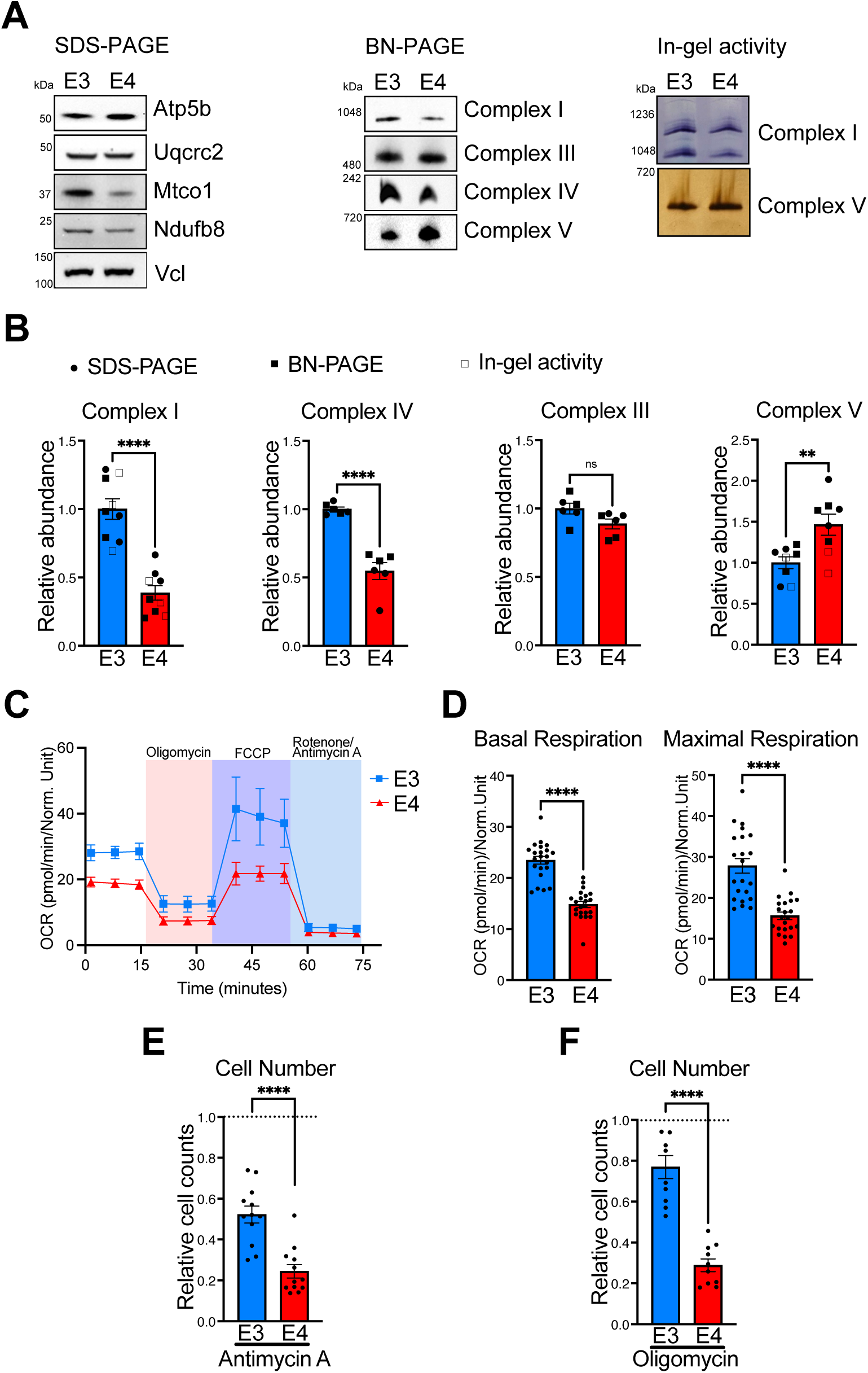
*APOE4* astrocytes display impaired respiration, and an extensively remodeled oxidative phosphorylation system. **A**) Whole cellular protein from *APOE3* (E3) and *APOE4* (E4) astrocytes fractionated by SDS-PAGE and immunoblotted for components of the oxidative phosphorylation system: complexes I (Ndufb8), III (Uqrc2), IV (Mtco1) and V (Atp5b) and the cytoskeletal protein vinculin indicates equal protein loading. Immunoblotting after blue native electrophoresis indicates the abundance of the intact OXPHS complexes, which are concordant with the individual denatured subunits. In-gel activity assays of complexes I and V after blue native electrophoresis. **B**) Overall abundance of complex I (p = 0.0001), complex IV (p = 0.0001), complex III and ATP synthase (p = 0.007) based on native and denaturing gel analysis. **C**) Representative profile of oxygen consumption rate (OCR) in astrocytes carrying the E3 or E4 variant. **D**) Basal and maximal respiration rate (OCR) in E3 and E4 astrocytes from n = 3 independent experiments, p = 0.0001. *APOE4* astrocytes are more sensitive to antimycin A and oligomycin than *APOE3* astrocytes. *APOE3* and *APOE4* astrocytes grown on 48 well plates were treated with **E**) 25 nM antimycin A, or **F**) 5 nM oligomycin for 48 hours, and cells numbers were derived from images of Hoechst-stained nuclei at 24 and 48 hours (n = 3 experiments and p = 0.0001 in both cases).

### Mitochondria of APOE4 astrocytes have an elevated membrane potential owing to reverse ATP synthase activity and low proton leak

High membrane cholesterol content has been previously linked to reduced proton leak across the inner mitochondrial membrane in isolated organelles (Baggetto et al., 1992). Concordantly, the mitochondrial proton leak in *APOE4* astrocytes was half that of the astrocytes carrying *APOE3* (**Figure 5A**), which suggests one function of cholesterol is to limit proton leak.

**Figure 5.**
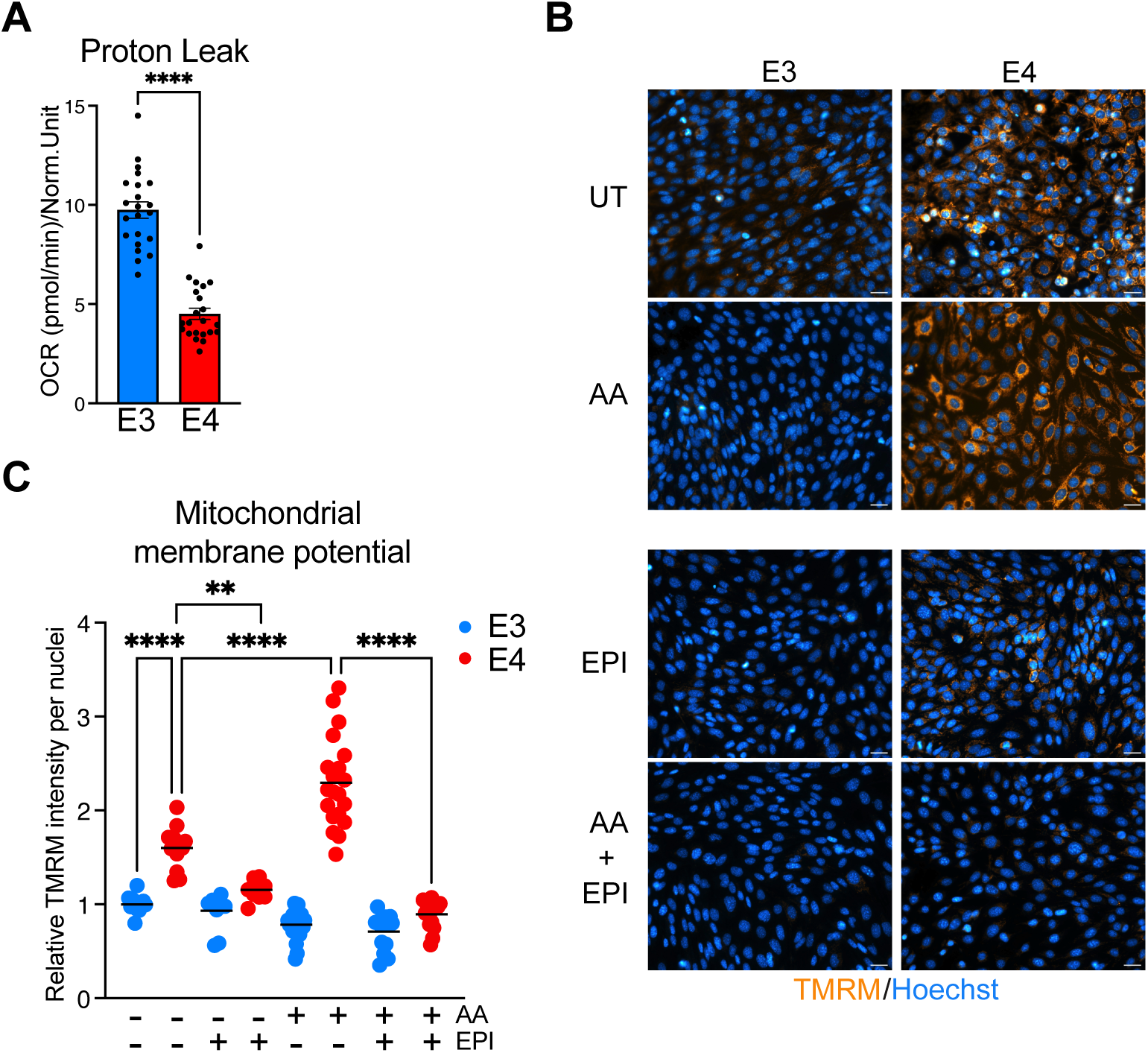
*APOE4* astrocytes have lower proton leak and higher reverse ATP synthase activity than *APOE3* cells. **A**) Flux analyzer determined proton leak, E3 versus E4, n = 3, p = 0.0001. **B**) Representative images of TMRM labeled *APOE3* and *APOE4* astrocytes incubated without or with 25 nM antimycin A (AA), and 100 μM epicatechin (EPI). Astrocytes were left untreated (UT) or treated for 16 hours with 100 nM antimycin A, with or without 100 μM epicatechin, and all were treated with 25 nM TMRM for 30 minutes on 96 well plates and microscopy images captured with identical settings. **C**) TMRM signal in APOE astrocytes with and without AA and EPI, n = 3 independent experiments, normalized to the signal of untreated APOE3 astrocytes. The APOE4 TMRM signal was 60% higher than that of *APOE3* cells without treatment, p = 0.0001; and 2.5-fold higher with AA, p = 0.0001. EPI decreased the *APOE4* TMRM signal 30%, p = 0.0011; and 2.6-fold, p = 0.0001, in the presence of AA, while having little effect on *APOE3* TMRM signal. One-way ANOVA non-parametric test was applied in all cases.

Although decreased proton leak will increase the proton gradient across the inner mitochondrial membrane, the low respiration capacity of the APOE4 mitochondria will result in a lower membrane potential, all other things being equal. In practice, the mitochondrial membrane potential was 60% higher in *APOE4* astrocytes compared to *APOE3* cells, as measured by the membrane potentiate dye, tetramethyl-rhodamine methyl ester (TMRM) (**Figure 5B, 5C**). Moreover, inhibition of respiratory complex III with antimycin A accentuated the difference in the mitochondrial membrane potential between *APOE4* and *APOE3* cells from 2 to 8-fold (**Figure 5B, 5C**). The proton ionophore FCCP collapsed the membrane potential in both cell lines, indicating that the TMRM signals were accurately reporting the mitochondrial proton gradient (**Figure S2**). This left ATP hydrolysis via ATP synthase operating in reverse (Yoshida et al., 2001) as the prime candidate for generating the high mitochondrial membrane potential of *APOE4* cells. As a test, we applied an inhibitor of reverse ATP synthase, epicatechin (Acin-Perez et al., 2023), with and without antimycin A. Exposure to 100 µM epicatechin for 16 hours decreased the mitochondrial membrane potential by 30% in *APOE4* astrocytes; whereas the decrease was 2.6-fold in the presence of 100 nM antimycin A compared to antimycin A alone (**Figure 5C**). The antimycin A-induced increase in mitochondrial membrane potential in APOE4 astrocytes was less marked at a dose of 25 nM for 48 hours; nevertheless, almost all the antimycin A-associated increase was abolished by 25 nM or 2.5 µM epicatechin (**Figure S3A, S3B**). Hence, the high membrane potential of APOE4 astrocytes, with and without antimycin A, is generated via reverse ATP synthase activity (**Figure 5B, 5C**).

### Inhibition of reverse ATP Synthase in APOE4 astrocytes attenuates cholesterol-induced cell death

Next we assessed possible functional consequences of the elevated reverse ATP synthase activity of APOE astrocytes. As inhibiting reverse ATP synthase with epicatechin proved no more toxic to *APOE4* than *APOE3* astrocytes (**Figure S3C**), we discarded the idea that *APOE4* astrocytes were dependent on a high mitochondrial membrane potential. Oppositely, we tested whether reverse ATP synthase activity was harmful, specifically whether it contributed to *APOE4* sensitivity to cholesterol overload. As well as reducing cell survival, cholesterol supplementation caused extensive cell clumping in APOE4 astrocytes (**Figure 6A**), which presumably reflects an excess of cholesterol in the plasma membrane. A low dose of 25 nM epicatechin halved cell clumping (**Figure 6A, 6B**) and increased the number *APOE4* astrocytes exposed to 10 µg/mL cholesterol for 48 hours 1.8-fold (**Figure 6C**). The greatest improvement in cell growth and survival was achieved with 2.5 µM epicatechin, which increased cholesterol-supplemented *APOE4* astrocyte numbers more than 2-fold (**Figure 6A, 6C**). Epicatechin also blunted antimycin A toxicity, although in this case the low dose of 25 nM epicatechin increased *APOE4* astrocyte numbers more than the higher dose of 2.5 µM epicatechin (1.7-fold versus 1.4-fold) (**Figure 6D**). Collectively, these data indicate that reverse ATP synthase activity contributes to maintenance of a high mitochondrial membrane potential, especially under conditions of severely impaired respiration. Inhibiting this maladaptive response is associated with increased tolerance to cholesterol and respiratory stress, manifesting as improved cell distribution, growth and survival.

**Figure 6.**
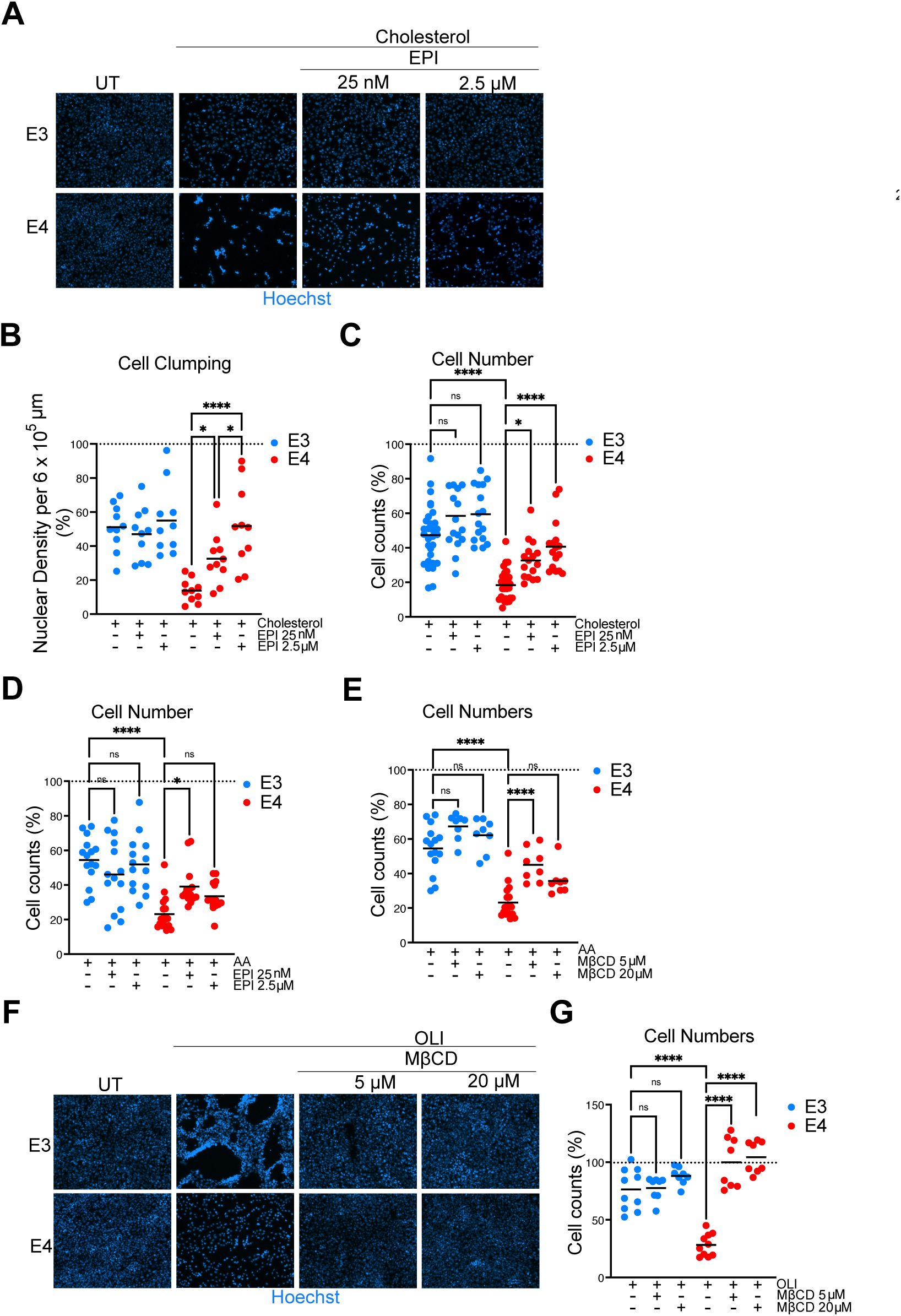
Epicatechin attenuates cholesterol toxicity in *APOE4* astrocytes, and cyclodextrin abrogates oligomycin-induced cell death. *APOE3* and *APOE* astrocytes were supplemented with 10 µg/mL cholesterol without or with epicatechin (EPI) for 48 hours and cells stained with Hoechst (**A**). Cell clumping (**B**) and cell number (**C**) were measured in n = 3 experiments. **D**) *APOE* astrocytes were treated for 48 hours with 25 nM antimycin A (AA), with and without 25 nM or 2.5 µM epicatechin, or left untreated, and stained with Hoechst and the cell numbers quantified in n = 3 experiments. **E**) As **D** except that methyl-β-cyclodextrin (MβCD) was used in place of epicatechin. **F**) *APOE3* and *APOE* astrocytes treated with 5 nM oligomycin (OLI), with and without 5 or 20 µM MβCD, or left untreated for 48 hours and stained with Hoechst, and cell numbers quantified from 3 independent experiments in panel **G**.

### Cholesterol sequestration counteracts ATP synthase dysfunction in APOE4 astrocytes

To further investigate cholesterol’s contribution to mitochondrial vulnerability in *APOE4* astrocytes, we tested whether reducing membrane cholesterol affected antimycin A sensitivity. We used methyl-β-cyclodextrin as it sequesters cholesterol from membranes and attenuates neurodegeneration in a mouse model of a cholesterol trafficking disorder (Davidson et al., 2009; Rosenbaum et al., 2010). While exposure to 25 nM antimycin A for 48 hours reduced the *APOE4* cell number to a quarter of untreated cells, co-treatment with 5 µM methyl-β-cyclodextrin increased cell number 1.6-fold, compared to antimycin A alone (**Figure 6E**). These findings suggest that the elevated intracellular cholesterol in *APOE4* astrocytes contributes directly to their sensitivity to the respiratory chain inhibitor.

We suspected that epicatechin’s ability to attenuate cholesterol and antimycin A toxicity was only one manifestation of ATP synthase dysfunction, and the sensitivity of *APOE4* astrocytes to oligomycin (**Figure 4F**) afforded us another means to test the hypothesis that elevated intracellular cholesterol provokes ATP synthase dysfunction. In this test, oligomycin was applied to the *APOE* astrocytes alone or in combination with methyl-β-cyclodextrin. While exposure to 5 nM oligomycin for 48 hours reduced the number of *APOE4* astrocytes to a quarter of untreated cells, co-treatment with 5 or 20 µM methyl-β-cyclodextrin abrogated the toxic effect of oligomycin, whereas the same treatments had a marginal effect on *APOE3* cell number (**Figure 6F, 6G**). Thus, we conclude that the ATP synthase dysfunction of *APOE4* astrocytes, that sensitizes them to oligomycin, stems from the elevated intracellular cholesterol.

Although we applied low doses of methyl-β-cyclodextrin in the micromolar range, small changes in cholesterol can produce marked changes in GPCR signaling, RTK activation, ion channels, endocytosis and cytoskeleton-membrane coupling, any of which might propagate the modest change in total cellular cholesterol to ATP synthase (Menon et al., 2024; Ward et al., 2026; Sun et al., 2007). That the cyclodextrin-induced changes in the mitochondria were subtle was supported by the facts that neither dose improved the cristae or the respiratory chain complexes in the *APOE4* astrocytes (**Figure S4**). Therefore, we next sought to achieve a more substantial decrease in intracellular cholesterol in the *APOE4* astrocytes.

### A cholesterol-lowering nutrient regime restores the cristae in APOE4 astrocytes

Ketone bodies reduce the high mortality associated with atherosclerosis in *ApoE*-deficient mice suggesting they mitigate the cholesterol perturbations caused by the lack of ApoE (Tomita et al., 2023). To create a similar nutrient status, astrocytes were cultured for 16 hours in a medium lacking glucose and pyruvate and supplemented with 0.6 mM β-hydroxybutyrate. Switching to this nutrient-restricted medium resulted in a more than 2-fold decrease in intracellular cholesterol, evidenced by BODIPY-cholesterol cell labeling (**Figure S5**) (Maekawa, 2017; Marks et al., 2008). Oppositely, pharmacological inhibition of cholesterol trafficking with U18666A increased BODIPY-cholesterol 5-fold in *APOE4* astrocytes, and more than an order of magnitude in *APOE3* cells (**Figure S5**).

Interpolating from these results, we speculated that *APOE4* astrocytes would be more tolerant of the cholesterol-lowering nutrient regime than *APOE3* astrocytes. In tests with culture medium, lacking glucose and pyruvate, with or without 0.6 mM β-hydroxybutyrate, *APOE4* astrocyte number was more than twice that of *APOE3* cells after 48 hours (**Figure 7A, 7B, and Figure S6**). Coupled with the opposite effect of cholesterol supplementation on the growth and viability of the two *APOE* variants (**Figure 1I**), these data suggest that the APOE4 alters cell cholesterol homeostasis in ways that drastically affect astrocyte sensitivity to cholesterol fluctuations - increasing vulnerability to exogenous cholesterol while mitigating low cholesterol levels.

**Figure 7.**
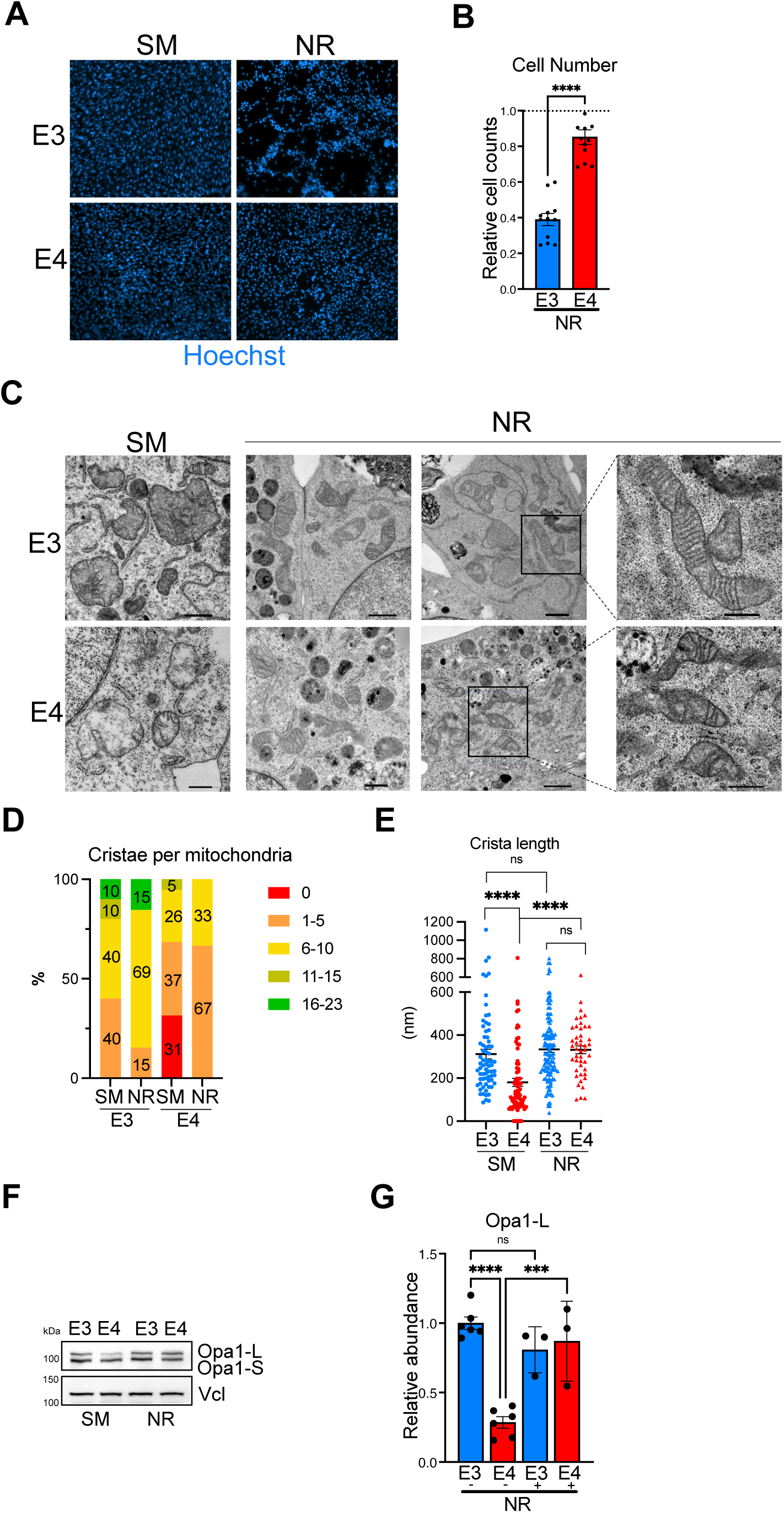
A cholesterol-lowering nutrient regime restores mitochondrial cristae in *APOE4* astrocytes. *APOE3* (E3) and *APOE4* (E4) astrocytes were cultured standard medium (SM) or in nutrient restricted medium (NR) lacking glucose and pyruvate with 0.6 mM β-hydroxybutyrate for 48 hours. **A)** Representative images of Hoechst-stained cells. **B)** Cell number after 48 hours of NR, relative to the same cells grown in parallel on SM, the latter set as 1 and indicated by a broken horizontal line. N = 3, p = 0.0001. **C)** Representative electron micrographs of astrocytes carrying *APOE3* (E3) or *APOE*4 (E4) variants grown on SM or NR for 16 hours. Scale bars from left to right = 1 µm, 1 µm, 1 µm and 500 nm. **D, E**) Charts indicating the difference in cristae number and length between mitochondria of *APOE3* and *APOE4* astrocytes with and without nutrient restriction. **F**) A representative immunoblot of Opa1 beside **G**) a chart of the abundance of the long and short isoforms of Opa1 in *APOE3* and *APOE4* astrocytes grown on SM or NR, from n = 3-6 experiments; p = 0.0001 between E3 and E4 in SM, and p = 0.0005 in E4 SM versus NR.

To assess the impact of the cholesterol lowering nutrient regime on the mitochondrial abnormalities associated with *APOE4* astrocytes, we analyzed the mitochondrial ultrastructure. Transmission electron microscopy of nutrient-restricted *APOE4* astrocytes revealed increased cristae length and number to the level of untreated *APOE3* astrocytes (**Figure 7C-7E**). Cristae structure was restored across the entire cell and mitochondrial population, evidenced by an increase in Opa1L to the level of *APOE3* astrocytes (**Figure 7F**). As the nutrient-restricted medium lowers cholesterol levels (**Figure S5**) and restores the architecture of the inner mitochondrial membrane (**Figure 7**), we infer that the disruption of the mitochondrial membranes is reversed owing to the lowering of intracellular cholesterol.

The increase in cristae number and length increases the surface area of the inner mitochondrial membrane, offering more space to accommodate respiratory chain complexes. Concordantly, there was a significant and greater than 2-fold increase in the abundance of respiratory complex IV in nutrient-restricted *APOE4* astrocytes (**Figure S7A, S7B**). Although respiratory complex I abundance was more variable and did not achieve statistical significance, it increased approximately 1.5-fold when *APOE4* astrocytes were switched to the β-hydroxybutyrate containing/nutrient restricted medium (**Figure S7C**). Finally, Atad3 expression increased in response to the cholesterol-lowering nutrient regime (**Figure S7D**) that restored the cristae (**Figure 7**), once again linking the triad of Atad3, cristae and cholesterol (Muñoz-Oreja et al., 2024) (Peralta et al., 2018).

## DISCUSSION

This study reveals a direct, mechanistic connection between elevated intracellular cholesterol, cristae collapse, and ATP synthase dysfunction that threatens cell viability. In *APOE4* astrocytes, elevated cholesterol coincides with reductions in cristae number and length and impaired oxidative phosphorylation, including increased reverse ATP synthase activity (**Figures 2–5**). These abnormalities are largely reversible: nutrient restriction lowers intracellular cholesterol and restores cristae architecture, while pharmacological interventions targeting either ATP synthase or cholesterol mitigate the lethal effects of the other (**Figures 6 and 7**). Thus, the inner mitochondrial membrane, and ATP synthase embedded within it, emerge as principal sensors and effectors of cellular cholesterol imbalance.

### A structural-functional model linking cholesterol, cristae, and ATP synthase

Integrating our current study with earlier findings (Peralta et al., 2018; Desai et al., 2017; Gunning et al., 2020; Kiesel et al., 2025; Muñoz-Oreja et al., 2024), we propose a model in which a tightly constrained cholesterol content is required for correct cristae formation and positioning of ATP synthase dimers for optimal ATP synthesis (**Figures 8 and S8**). Excess cholesterol stiffens biological membranes altering local curvature (Doole et al., 2022), which in the context of the inner mitochondrial membrane would be expected to disrupt the ordered rows of ATP synthase dimers that shape and stabilize cristae ridges (Blum et al., 2019; Buzzard et al., 2024; Davies et al., 2012; Paumard et al., 2002). Because ATP synthase dimers are juxtaposed at a precise angle, even modest increases in cholesterol and subtle changes in cristae curvature could impair ATP synthase function, consistent with the proposition that ATP synthase performance is a function of membrane positioning (Strauss et al., 2008). In *APOE4* astrocytes, the disruption of cristae architecture shifts ATP synthase toward reverse activity, and the heightened oligomycin toxicity demonstrates the enzyme’s central role in cell vulnerability under conditions of cholesterol excess.

**Figure 8.**
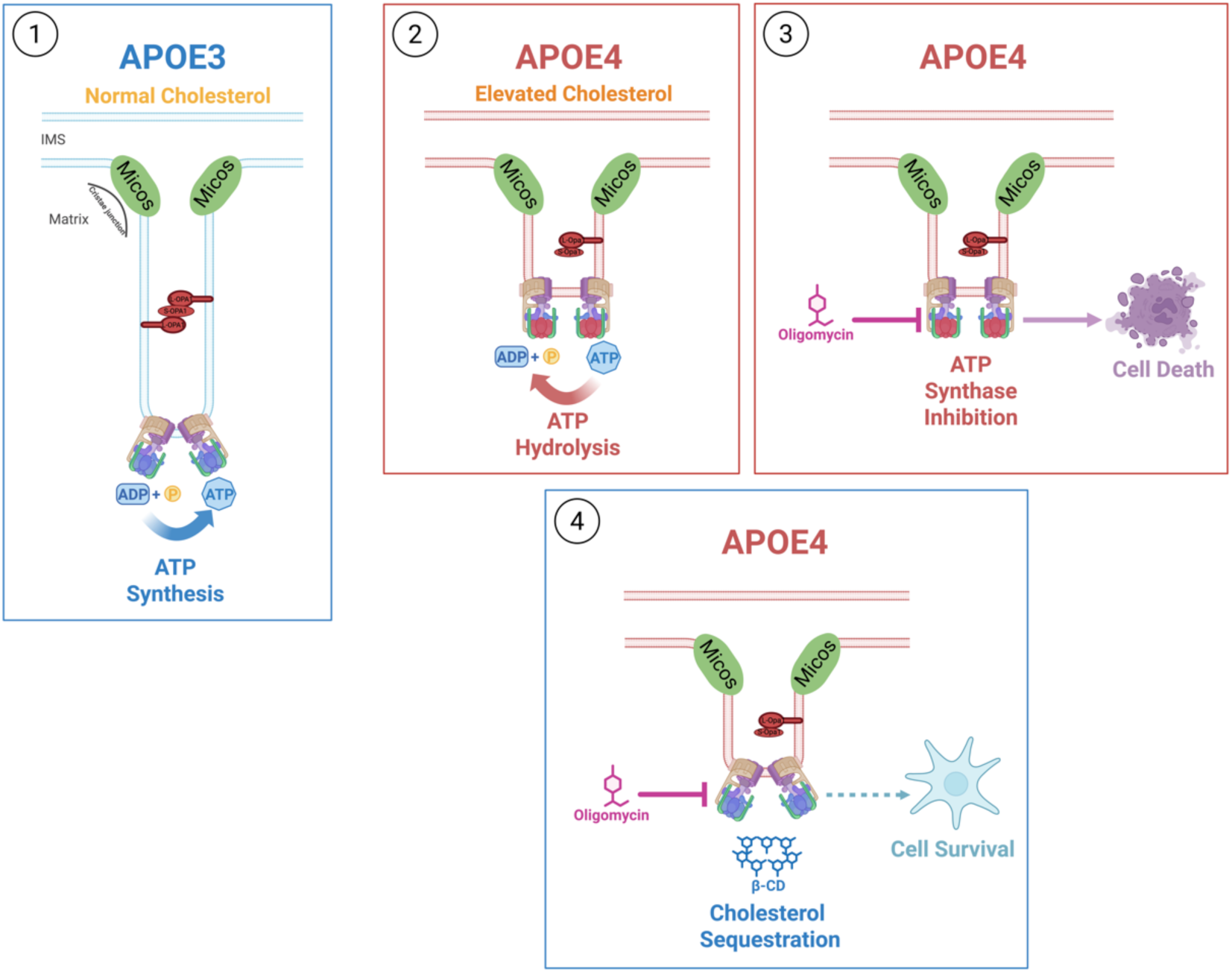
Cholesterol is a key determinant of cristae architecture and ATP synthase function. **1)** Cristae and ATP synthase organization when cholesterol homeostasis is functioning well, as per *APOE3* astrocytes. **2)** Elevated intracellular cholesterol in *APOE4* astrocytes disrupts the inner mitochondrial membrane, manifesting as truncated and scarce cristae with reduced Opa1-L and altered ATP synthase. **3)** Oligomycin is extremely toxic to APOE4 astrocytes, but **4)** is rescued by cholesterol sequestration with cyclodextrin (β-CD) demonstrating causality. Although the precise arrangement of ATP synthase in conditions of elevated cholesterol is not known, the maintenance of holoenzymes that exceed in number those of APOE3 astrocytes (Figure 4A) suggests the defect is subtle, and so it is represented in the model as a change in the dimer angle.

The parallels between APOE4 (this report) and Atad3 loss (Peralta et al., 2018) are striking. Both feature perturbed cholesterol homeostasis, cristae disruption and the accumulation of cristae junction complexes. These findings support our earlier proposal that ATAD3 introduces or organizes cholesterol in mitochondrial membranes (Muñoz-Oreja et al., 2024), with the new insight that mitochondrial cholesterol, together with ATAD3, plays a critical role in cristae maintenance.

### ATP synthase as a nexus for aberrant cholesterol homeostasis

A key question is the relative importance of mitochondrial changes resulting from elevated intracellular cholesterol versus alterations to other organelles. Two lines of evidence calibrate the mitochondrial contribution. First, and most compellingly, cholesterol sequestration with cyclodextrin abrogates oligomycin toxicity selectively in *APOE4* astrocytes (**Figure 6F, 6G**), directly demonstrating that elevated cholesterol is the proximate cause of ATP synthase malfunction rather than a correlative finding. Second, inhibition of reverse ATP synthase with epicatechin attenuates cholesterol-induced cell death (**Figure 6A-6C**), confirming that reverse ATP synthase activity is a meaningful downstream effector of cholesterol excess, albeit one that accounts for only part of the cellular vulnerability. Together, these findings define a causal axis - from cholesterol accumulation through cristae collapse to ATP synthase dysfunction and cell vulnerability - that operates independently of the sterol’s effects elsewhere in the cell.

### Disease relevance and translational opportunities

The APOE4 context links our mechanistic findings to human disease, as *APOE4* is the strongest common genetic risk factor for Alzheimer’s disease, and *APOE4* homozygosity constitutes a distinct genetic form accounting for ~15% of cases (Fortea et al., 2024). Separately, elevated LDL cholesterol is a population-level risk factor for both atherosclerosis and dementia (Livingston et al., 2024). Our results implicate cholesterol-mediated cristae remodeling and ATP synthase disruption as a unifying mechanism that could operate across cholesterol disorders from atherosclerosis to neurodegeneration (Duan et al., 2022; TCW et al., 2022). That nutrient restriction restores cristae (**Figure 7**) and cyclodextrin abrogates oligomycin toxicity (**Figure 6**) suggest testable therapeutic strategies for cholesterol-related disorders based on cholesterol restriction via dietary or pharmacological control. Moreover, ATP synthase function and cristae ultrastructure emerge as candidate biomarkers for patient stratification and treatment monitoring. That said, the threat to the mitochondrial inner membrane and ATP synthase posed by the elevated intracellular cholesterol of *APOE4* astrocytes may elicit mitigating responses. The apparent discrepancy between the impaired respiration we report in proliferating *APOE4* astrocytes and the elevated respiration associated with senescent *APOE4* astrocytes (Caceres-Palomo et al., 2025) can, we propose, be reconciled and is itself informative. The proliferative imperative of immortalised cells brings into sharp relief the mitochondrial vulnerability imposed by cholesterol excess - collapsed cristae, impaired oxidative phosphorylation, and pathological reverse ATP synthase activity. Senescence, by contrast, has been shown to promote cristae elaboration and a shift toward fatty acid oxidation (Yamauchi et al., 2024) - a metabolic state that is almost the opposite of the one we describe. This striking symmetry supports the interpretation that senescence is induced as an adaptive response to cholesterol-driven mitochondrial stress in primary human *APOE4* astrocytes, with the restructured cristae and altered substrate utilisation of the senescent cell directly counteracting the vulnerabilities uncovered here in immortalized astrocytes for whom senescence is not an option.

A further indication that cholesterol homeostasis is finely balanced comes from the observation that *APOE4* astrocytes tolerate nutrient restriction better than their *APOE3* counterparts. This suggests that elevated intracellular cholesterol, whilst detrimental under nutrient-rich conditions, may confer resilience under nutrient restriction. Since nutrient restriction lowers intracellular cholesterol, the larger cholesterol pool of *APOE4* astrocytes sustains them longer under these conditions, whereas *APOE3* astrocytes, starting from a lower baseline, reach a critical threshold sooner.

### Limitations and future directions

Cholesterol affects membrane rigidity over considerable distances at the molecular scale, and so it need not be intimately associated with cristae ridges or ATP synthase to exert the effects we observe. Determining the precise distribution of cholesterol within mitochondrial subdomains will be a major challenge, as even the most accessible cholesterol sensors focus chiefly on the cholesterol-rich plasma membrane, and none gives information on individual microdomains. Cryo-electron tomography and super-resolution imaging may ultimately reveal how elevated cholesterol alters dimer angles or row continuity of ATP synthase (Buzzard et al., 2024; Blum et al., 2019), and how this integrates with OPA1/MICOS dynamics (Del Dotto et al., 2017; Hu et al., 2020).

The comparison between immortalised murine astrocytes, in which the proliferative imperative prevents senescence, and primary human *APOE4* astrocytes, which can mount an adaptive senescent response, illustrates the value of complementary cell models in uncovering the full spectrum of cholesterol-driven mitochondrial vulnerability. Whether the findings extend to other cholesterol-handling cell types such as neurons or hepatocytes remains to be established. More broadly, our finding that withdrawal of glucose and pyruvate restores the cristae demonstrates that the metabolic environment profoundly influences mitochondrial outcomes in *APOE4* astrocytes, indicating that the phenotypes we describe are not strictly cell-autonomous. *In vivo*, that environment is shaped by neurons, microglia, and other cell types, whose metabolic and signalling outputs affect astrocyte function. Reciprocally, the cholesterol that astrocytes package and supply to neurons and other cell types means that APOE4-driven dyshomeostasis in the astrocyte may disturb the broader cholesterol economy of the brain, creating a self-reinforcing cycle of lipid imbalance and mitochondrial vulnerability.

## Conclusion

We identify cholesterol-driven cristae disruption as a proximate cause of ATP synthase dysfunction and cell vulnerability in *APOE4* astrocytes, positioning mitochondrial cholesterol as a critical determinant of cristae architecture and bioenergetic integrity. The bidirectional rescue experiments reported here - in which targeting cholesterol abrogates ATP synthase vulnerability and targeting ATP synthase attenuates cholesterol toxicity - establish causality and call for systematic assessment and modulation of cristae morphology, ATP synthase behaviour, and mitochondrial cholesterol across the spectrum of cholesterol-related and mitochondrial diseases.

## METHODS

### Cell Culture

Immortalized astrocyte cell lines derived from knock-in mice expressing human *APOE3* or *APOE4* were as previously described (Morikawa et al., 2005). Astrocytes were maintained in Dulbecco’s Modified Eagle Medium (DMEM) containing 25 mM glucose and 1 mM sodium pyruvate, supplemented with 10% heat-inactivated fetal bovine serum, 2 mM GlutaMAX, and penicillin-streptomycin (100 U/mL penicillin and 100 µg/mL streptomycin; (Gibco). Astrocytes were cultured at 37°C in a humidified incubator with 5% CO_2_. The cell lines were routinely tested and confirmed to be mycoplasma-free, using the Venor®Gem Classic Mycoplasma PCR Detection Kit (Minerva Biolabs).

### Mitochondrial respiration

Astrocytes 0were counted using a TC20 Automated Cell Counter (BioRad) and seeded into XF Pro M Cell Culture Microplates (Agilent Technologies) at a density of 8 × 10^3^ cells per well in 80 µl of standard culture medium. Cells were cultured for 24 h prior to analysis. For mitochondrial stress tests, the culture medium was replaced with 180 µL of Seahorse XF DMEM medium (pH 7.4; Agilent Technologies). Cells were incubated for 1 h at 37°C in a non-CO_2_, humidified incubator to allow temperature and pH equilibration. Oxygen consumption rate (OCR) was measured using a Seahorse XF Pro Analyzer (Agilent Technologies). Basal respiration was recorded over three measurement cycles, followed by sequential injections of mitochondrial inhibitors. Oligomycin was injected to a final concentration of 2.5 µM, and OCR was measured over three cycles to assess ATP-linked respiration. FCCP (carbonyl cyanide 4-(trifluoromethoxy) phenylhydrazone) was then injected to a final concentration of 2 µM, followed by three measurements to determine maximal respiratory capacity. Finally, rotenone and antimycin A were injected simultaneously to final concentrations of 0.5 µM each, and OCR was measured over three cycles to quantify non-mitochondrial oxygen consumption. Oligomycin A, FCCP, rotenone, and antimycin A were purchased from Merck (Sigma-Aldrich). All compounds were prepared as 10 mM stock solutions in ethanol and stored at −20 °C.

### Survival assays

Astrocyte survival was assessed using live-cell imaging. Astrocytes were counted using a TC20 Automated Cell Counter (BioRad) and seeded at a density of 1 × 10^4^ cells per well in 48-well cell culture plates (Costar). After 24 h, cells were treated with the following compounds at the indicated final concentrations: cholesterol (10 µg/mL from the Abcam kit; or its equivalent 3 x Gibco cholesterol supplement Cat # 12531018), 25 nM antimycin A (Sigma-Aldrich), 5 nM oligomycin (Sigma-Aldrich), 5 µM or 20 µM β-cyclodextrin (MedChemExpress), and 25 nM or 2.5 µM epicatechin (MedChemExpress). At either 24 h or 48 h post-treatment, depending on the experimental design, astrocytes were stained with 1:6,000 Hoechst 34580 (Invitrogen) for 15 min at 37°C in 5% CO_2_. Cells were subsequently washed three times with Dulbecco’s phosphate-buffered saline (DPBS; Gibco) and maintained in FluoroBrite™ Live Cell Imaging Medium (Gibco). Live-cell images were acquired using an LSM 900 confocal microscope with incorporated CO_2_ and temperature control modules (Zeiss).

### Mitochondrial Membrane Potential (ΔΨm)

Astrocytes were counted using a TC20 Automated Cell Counter (BioRad) and seeded at a density of 8 × 10^3^ cells per well in 96-well microplates (ibidi GmbH). After 24 h, cells were treated without and with antimycin A (25 nM or 100 nM) and/or epicatechin (25 nM or 100 µM; MedChemExpress) for 16 h at 37°C in 5% CO_2._ Mitochondrial membrane potential (ΔΨm) was assessed using 25 nM tetramethylrhodamine methyl ester (TMRM) (Invitrogen) for 30 min at 37°C, followed by nuclear staining with a 1 in 6 000 dilution of Hoechst 34580 (Invitrogen) for 15 min. Cells were washed three times with DPBS (Gibco) and maintained in FluoroBrite™ Live Cell Imaging Medium(Gibco). Live-cell images were acquired using an LSM 900 confocal microscope with incorporated CO_2_ and temperature control modules (Zeiss).

### Transmission Electron Microscopy

Prior to imaging, astrocytes were cultured for 24 h in either standard medium or medium in which glucose and pyruvate were replaced with 0.6 mM β-hydroxybutyrate. Cells grown on eight-well Lab-Tek®II chamber slides^TM^ (Nunc) were washed with 0.1 M phosphate buffer (NaH_2_PO_4_·H_2_O and Na_2_HPO_4_) and fixed with 3% glutaraldehyde for 10 min at 37°C followed by 2 h at room temperature. Samples were washed five times with phosphate buffer (5 min per wash), air-dried, and subsequently processed for electron microscopy at the Centro de Investigación Príncipe Felipe (Valencia, Spain). Ultrathin sections were imaged using a 200-kV high-resolution TECNAI G2 20 TWIN transmission electron microscope, as previously described (Muñoz-Oreja et al., 2024).

### Immunocytochemistry (ICC)

8 × 10^3^ astrocytes per well were seeded on 96-well microplates (Ibidi) and after treatment fixed with 4% paraformaldehyde (PFA; Electron Microscopy Sciences) for 20 min at room temperature and washed twice with dPBS (Gibco). Cells were permeabilized with 0.3% Triton X-100 PBS (PBST) for 15 mins at room temperature and subsequently blocked in 0.1% Triton X-100 PBS (PBST) containing 10% donkey (GeneTex) or goat serum (Sigma), depending on the secondary antibodies used, for 1 h at room temperature. Primary antibodies were applied in PBST overnight at 4°C: anti-Atad3 and anti-Mic60 (Proteintech). After three 5-min washes with PBST, cells were incubated with secondary antibodies, Alexa Fluor^TM^ 488/555 goat/donkey anti-mouse IgG (H+L) and Alexa Fluor^TM^ 488/555 goat/donkey anti-rabbit IgG (H+L) from Invitrogen diluted 1:450 in 10% serum/PBST for 1 h at room temperature. Cells were washed two times with PBST, and once with PBS and dH_2_O, and nuclei were counterstained using mounting medium with DAPI (ibidi GmbH), as previously (Muñoz-Oreja et al., 2024).

### Gm1 labeling

Following PFA fixation, cells were protected from light and incubated overnight at 4°C with 1 µg/ml FITC-conjugated Cholera Toxin B Subunit (Sigma) in PBS containing 1% BSA, with or without 2% saponin (0.5% w/v in PBS; Thermo Scientific). After three 5-min washes with DPBS, cells were incubated with secondary antibodies (Alexa Fluor^TM^ 488 goat anti-mouse IgG (H+L), Alexa Fluor^TM^ 555 goat anti-rabbit IgG (H+L); Invitrogen), at 1:2000 dilution in DPBS for 1 h at room temperature. Cells were washed three times with DPBS and mounted with DAPI-containing medium (ibidi GmbH).

### Bodipy-cholesterol labeling

After incubating cells with 1 µM BODIPY™-Cholesterol (MedChemExpress, HY-125746) for 4 hours, cells were washed with DPBS, FluoroBrite™ Live Cell Imaging Medium (Gibco) before adding Hoechst (Invitrogen, 1:5000) and proceeding to image acquisition.

### Image capture and analysis

Fluorescent images were acquired with a LSM 900 Zeiss confocal microscope with incorporated CO_2_ and temperature control modules (Zeiss). Laser power, gain and offset parameters were constant for each experiment; any subsequent adjustments to contrast and brilliance were applied equally to all images. The image analysis was performed using Fiji ImageJ software and GraphPad (Prism) was used to quantify and represent the data as charts.

### SDS-PAGE

Astrocytes were detached from the flasks with trypsin (0.4% w/v solution, Gibco), reaction inhibited with DPBS containing 10% FBS, and pelleted by centrifugation at 500 × g for 5 min at room temperature. Pellets resuspended in DPBS underwent microcentrifugation for 500 × g for 5 min at 4°C, and lysed on ice with RIPA buffer (150 mM NaCl, 1% Triton X-100, 0.5% sodium deoxycholate, 0.1% SDS, 50 mM Tris-HCl, pH 8.0; Sigma-Aldrich) supplemented with Halt™ Protease and Phosphatase Inhibitor Cocktail (Thermo Scientific) for 30 min. Lysates were centrifuged at 20,000 × g for 20 min at 4°C, and protein concentration was determined using the DC Protein Assay Kit (Bio-Rad). Proteins (10 µg/well) were supplemented with Laemmli buffer containing DTT (Invitrogen) and resolved on 8%, 10%, and 12% Mini-PROTEAN TGX™ precast gels (Bio-Rad) using Tris/Glycine/SDS running buffer at 150 V for 70 min.

### BN-PAGE

Astrocytes pellets were resuspended in 1 x protein solubilizing solution (PSS), which was made by 1 M 6-aminocaproic acid, 50 mM bis-tris (Sigma-Aldrich); protein concentration was measured using the DC Protein Assay Kit. 100 µg sample were pelleted at 12,000 × g for 10 min, resuspended in 5% (w/v) digitonin stock solution (Thermo Scientific) was added to a final concentration of 2g/g or 4g/g digitonin:protein ratio with the protease inhibitor described above, incubated on ice for 5 min, and centrifuged at 20,000 × g for 20 min at 4°C. Supernatant was resuspended with 5% Serva Blue G (Sigma-Aldrich), and 20 µg per well was loaded onto 3-12% gradient native polyacrylamide gels (Invitrogen). Gels were run with cathode buffer A (50 mM tricine, 15 mM Bis-Tris, 0.02% Serva Blue G, pH 7.0; Sigma-Aldrich) and anode buffer (50 mM Bis-Tris, pH 7.0; Sigma-Aldrich) at 100 V for 15 min, followed by constant current (4 mA) for 2.5 h.

### Clear Native PAGE

Pellets were resuspended in PSS containing 0.5% n-dodecyl-β-D-maltoside (Sigma-Aldrich) and centrifuged at 4,700 × g for 5 min at 4°C. Protein concentration of the supernatants was determined using the DC Protein Assay Kit. Samples (20 µg/well) were mixed with 1/10 volume 50% glycerol and 1/10 volume 0.1% Ponceau S and separated on 3-12% native polyacrylamide gels. Gels were run with cathode buffer C (50 mM tricine, 15 mM Bis-Tris, 0.05% Triton X-100, 0.05% deoxycholate; Sigma-Aldrich) and anode buffer (50 mM Bis-Tris, pH 7.0; Sigma-Aldrich) at 100 V for 15 min, followed by constant current (4 mA) for 2.5 h (Aref et al., 2025).

### Immunoblotting

For denatured proteins electro-transfer to low-fluorescence PVDF membranes (Thermo Scientific) at 25 V, 90 mA for 16 h used Towbin buffer (25 mM Tris, 192 mM glycine, 20% methanol; Sigma-Aldrich). Membranes were blocked with a solution of 5% skimmed milk powder (Sigma-Aldrich) with 0.1% Tween 20 (PanReac AppliChem) in DPBS for 1 hour at room temperature and incubated overnight at 4°C with primary antibodies in the same buffer. After three 10-minute washes with 0.1% Tween-20/DPBS, membranes were incubated for 1 h at room temperature with 1:5000 dilutions of the appropriate secondary antibodies, HRP conjugated Goat Anti-Mouse IgG (H+L) and Goat Anti-Rabbit IgG (H+L), washed 3x with 0.1% Tween-20/DPBS, and developed using Clarity Western Peroxide and Luminol/Enhancer reagents (1:1, Bio-Rad). Signals were acquired with an iBright FL1500 Imaging System (Invitrogen) and quantified using ImageJ (FIJI). For proteins separated on native gels, transfer to PVDF membranes was in Towbin buffer at 100 V for 1.25 h. Membranes were blocked with 10% skimmed milk in DPBS for 1 h and incubated overnight at 4°C with primary antibodies in 0.3% Tween-20/PBS. After three 10-min washes, membranes were incubated with the appropriate secondary antibodies, HRP conjugated Goat Anti-Mouse IgG (H+L) and Goat Anti-Rabbit IgG (H+L) from Promega diluted in 1 in 5000, for 1 h at room temperature, washed, and developed as above. The type, source, and dilution of the other antibodies used in the study are listed in Table S1.

### *In-gel* activity assays

#### NADH Dehydrogenase (Complex I) Activity Staining

NADH dehydrogenase activity was assessed by in-gel staining following BN-PAGE. Gels were incubated in a staining solution comprising 2 mM Tris-HCl (pH 7.4), 2 mg NADH, and 50 mg nitroblue tetrazolium (Sigma-Aldrich) in a total volume of 20 mL. Incubation was performed at 37°C with gentle agitation until the enzymatic activity was sufficient to generate visible bands, at least in controls. The reaction was stopped by rinsing the gels thoroughly with double-distilled water (ddH_2_O), as previously described (Zerbetto et al., 1997).

#### ATP Synthase (Complex V) Activity Staining

ATPase activity was detected by in-gel following clear native PAGE. Gels were incubated at 37°C with gentle agitation in staining buffer consisting of 34 mM Tris, 270 mM glycine, 14 mM MgSO_4_, 0.2% (w/v) Pb (NO_3_)_2_, and 8 mM ATP, pH 7.8 (Sigma-Aldrich). To enhance staining, a 1.5% (w/v) ammonium sulfide (NH_4_)_2_S solution (Sigma-Aldrich) was added after initial incubation, and gels were gently agitated to ensure rapid and uniform coverage. After 10 seconds, gels were rinsed extensively with ddH_2_O to terminate the reaction, as previously described (Aref et al., 2025).

### Statistical Analysis

Statistical analyses were performed using GraphPad Prism (version 9.0; GraphPad Software). Data normality was assessed using the Shapiro-Wilk test. For comparisons between two independent groups, normally distributed data were analyzed using an unpaired, two-tailed Student’s *t*-test, whereas non-normally distributed data were analyzed using the Mann-Whitney *U* test. For comparisons involving more than two groups, one-way analysis of variance (ANOVA) was used, followed by appropriate post hoc tests as indicated in the figure legends. Statistical significance was defined as *P* ≤ 0.05. Significance levels are denoted as *P ≤ 0.05, **P ≤ 0.01, ***P ≤ 0.001, ****P ≤ 0.0001. Exact *P* values are reported in the figure legends. Unless otherwise stated, data are presented as mean ± standard error of the mean (SEM). Sample sizes (*n*), statistical tests used, and measures of variance are specified in the figure legends.

## Acknowledgements & Funding

The astrocyte lines were kindly provided by Prof. Mazhair Hasan, Basque Institute for Neuroscience, Bilbao, Spain. S.L is supported by the European Commission MitGEST (Mitochondrial Genome Stability), Marie Skłodowska-Curie Innovative Training Network (ITN). Grant Agreement No. 101073206. Horizon Europe Programme. M.M.O. was supported by a predoctoral fellowship from the University of the Basque Country (PIF18/317). A.L. and U.F.P. were recipients of pre-doctoral fellowships from the Basque Government (PRE_2019_1_0184 and PRE_2018_1_0253). A.L. was subsequently supported by the European Union project PMPPER24/00017_SEED-ALS. IJH gratefully acknowledges the support of the Spanish Ministry of Health (ISCIII: PI20/00096), and subsequently the Ministry of Science and Innovation (PID2023-151649NB-I00), as well as grants from the Basque Government Department of Health (Osasun Saila, Eusko Jaurlaritzako, grants 2021111070; 2022333050; 2018111043; 2018222031). AS was the recipient of a Miriam Marks Senior Fellowship, Brain Research UK (2021-2026) and is also supported by the Lily Foundation. AS and IJH are supported jointly by UK Medical Research Council grant MR/X002365/1.

