## Supplemental Information for "Elevated cholesterol in APOE4 astrocytes drives mitochondrial cristae collapse and ATP synthase dysfunction"

Seungtae Lee et al.

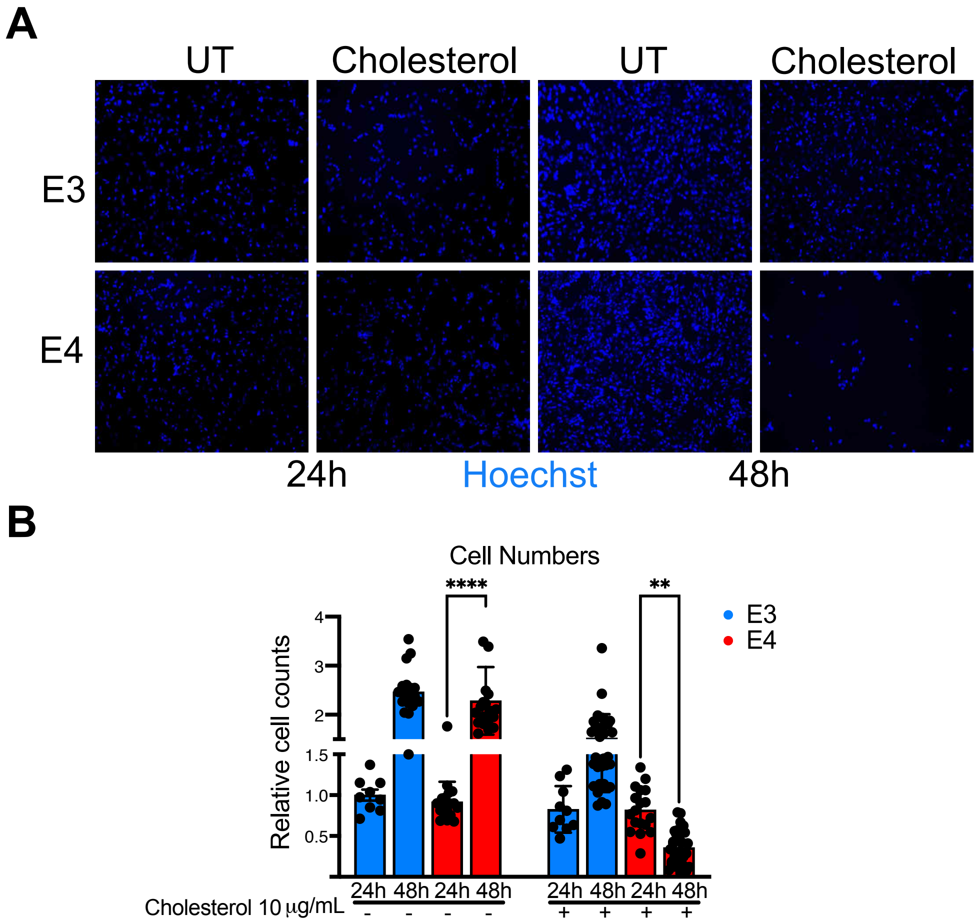


**Figure S1. Cholesterol supplementation causes astrocyte cell death not merely growth arrest. A)** hAPOE4 and hAPOE3 astrocytes grown on 48 well plates were treated without (UT) and with 10 µg/mL cholesterol for 48 hours and cells numbers were based on Hoechst stained nuclei at 24 and 48 hours. **B**) Quantification of three independent experiments; E4 cell number fell between 24 and 48 hours in the presence of 10 µg/mL cholesterol ($p = 0.004$).

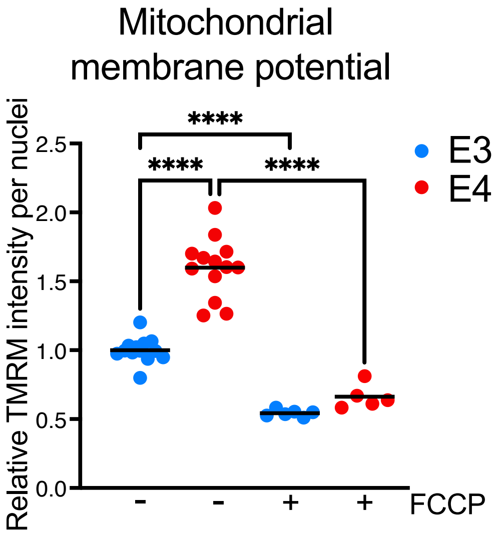


**Figure S2. The proton ionophore FCCP collapses the mitochondrial proton gradient in APOE3 and APOE4 astrocytes.** TMRM signal was quantified via ImageJ in APOE3 and APOE4 astrocytes incubated with and without 10 µM of the proton ionophore FCCP for 15 minutes, P = 0.0001.

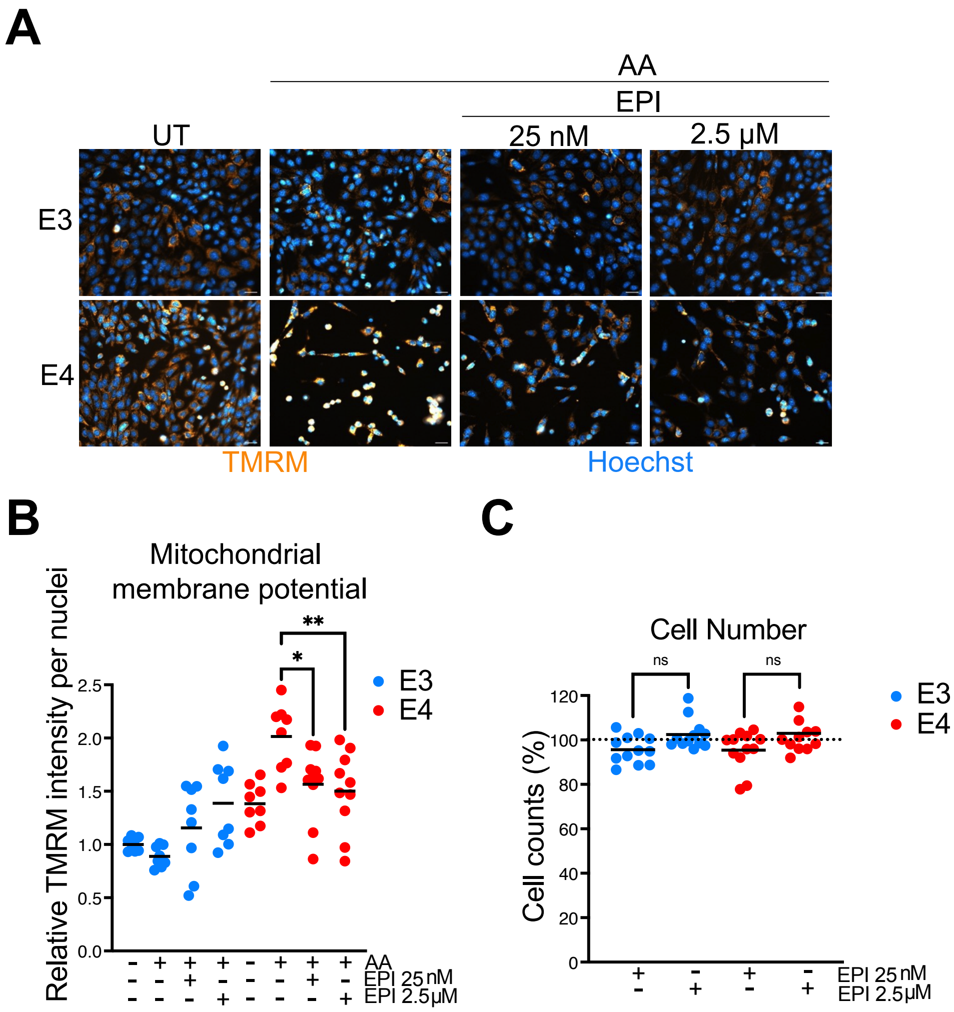


**Figure S3. 25 nM or 2.5 µM Epicatechin lower the mitochondrial membrane potential and are equally well tolerated by APOE3 and APOE4 astrocytes. A)** APOE3 and APOE4 astrocytes were left untreated or treated with 25 nM antimycin A (AA), with and without 25 nM or 2.5 µM epicatechin, for 48 hours and stained with Hoechst and TMRM, and **B**) the latter quantified in n = 3 experiments. **C)** APOE3 and APOE4 astrocytes were treated with and without 25 nM or 2.5 µM epicatechin, for 48 hours and stained with Hoechst. The cell number was quantified in n = 3 experiments and expressed relative to the number of cells receiving the vehicle in parallel.

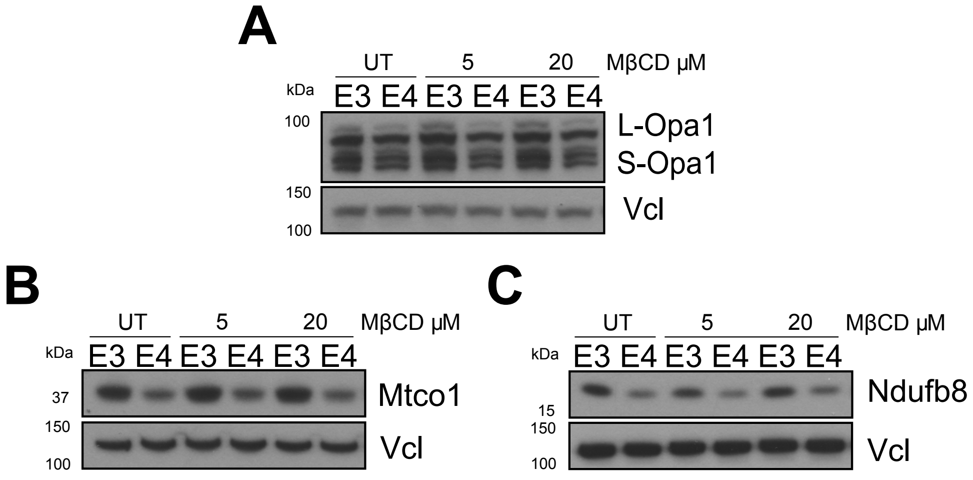


**Figure S4**. **Methyl-β-cyclodextrin does not rescue cristae structure or respiratory chain complex abundance in *APOE4* astrocytes. A)** Immunoblot analysis of the cristae-associated proteins Opa1 in APOE3 and APOE4 astrocytes treated with 5 or 20 µM MβCD for 24 hours, or left untreated, with vinculin (Vcl) as a loading control. **B)** Immunoblot of the complex IV subunit Mtco1 under the same treatment conditions, showing no restoration of complex IV levels. **C)** Immunoblot of the complex I subunit Ndufb8 under the same conditions, indicating no recovery of complex I abundance.

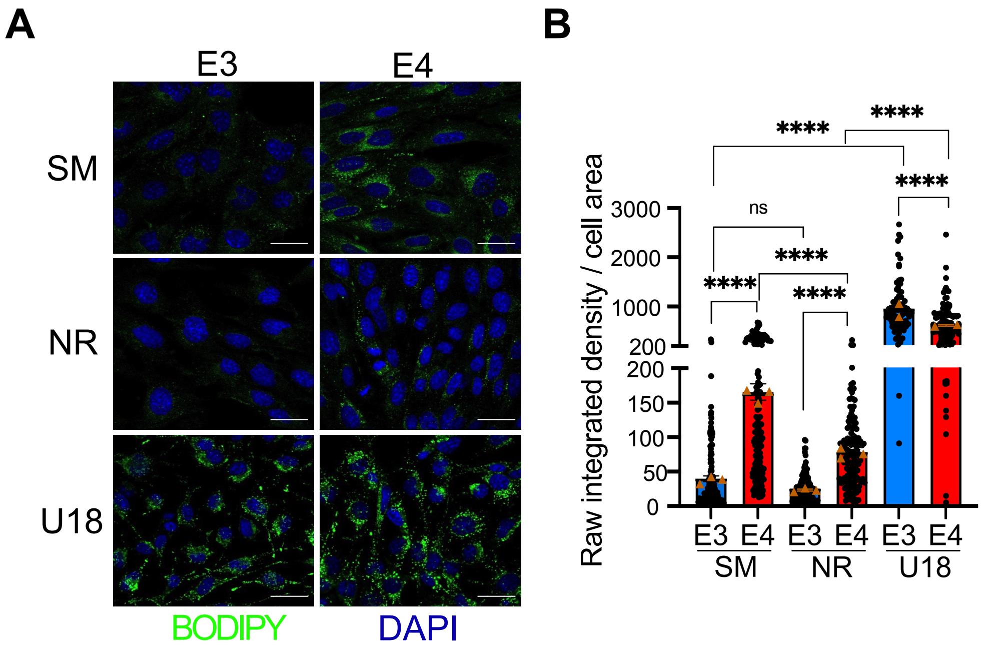


**Figure S5. Nutrient restriction lowers the intra-cholesterol level, whereas U18666A raises it.** Murine astrocytes were cultured in medium containing 25 mM glucose and 1 mM pyruvate (standard medium, SM), or in a nutrient restricted (NR) medium containing 0.6 mM β-hydroxybutyrate in place of glucose and pyruvate, or in SM with 2.5 µM U18666A, for 24 hours. **A)** Cells were live-stained with BODPY-cholesterol, both green and the nuclei counterstained blue with DAPI. Scale bar = 50 µm. **B)** Quantification of BODIPY-cholesterol signal (measured as raw integrated density per total area of the cell), where each point represents a cell and each colour a different cell line [>100 cells per line, n = 3 independent experiments, except U18666A where n = 2].

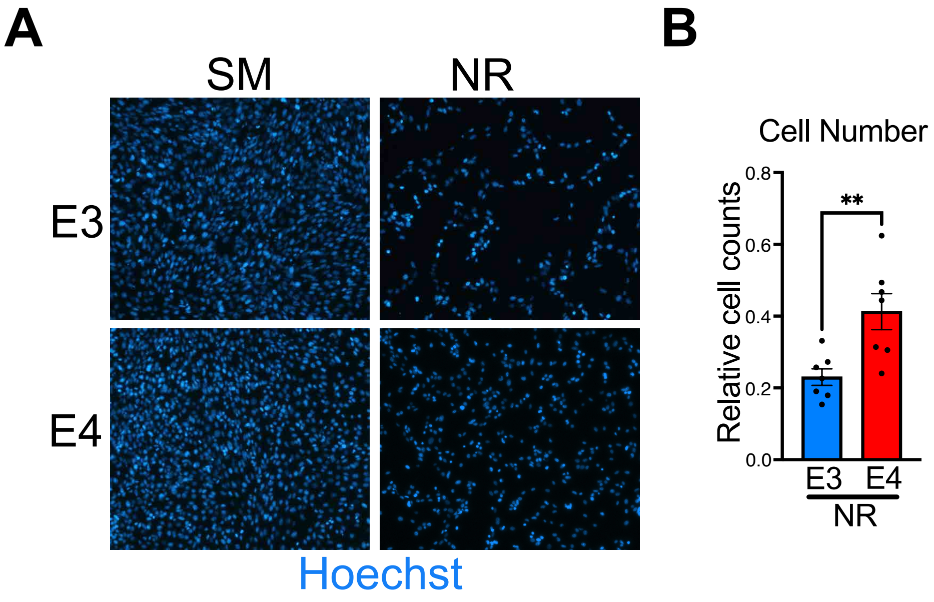


**Figure S6. APOE4 astrocytes tolerate a nutrient restricted medium lacking glucose and pyruvate better than APOE3 astrocytes.** APOE3 and APOE4 astrocytes were cultured in standard medium (SM) or in a nutrient restricted medium lacking glucose and pyruvate (NR) for 48 hours. **A)** Representative images of Hoechst-stained cells. **B)** Cell number after 48 hours of NR relative to the same cells grown on SM in parallel; the latter set as 1 and indicated by a broken horizontal line, n = 3 experiments, p =0.006.

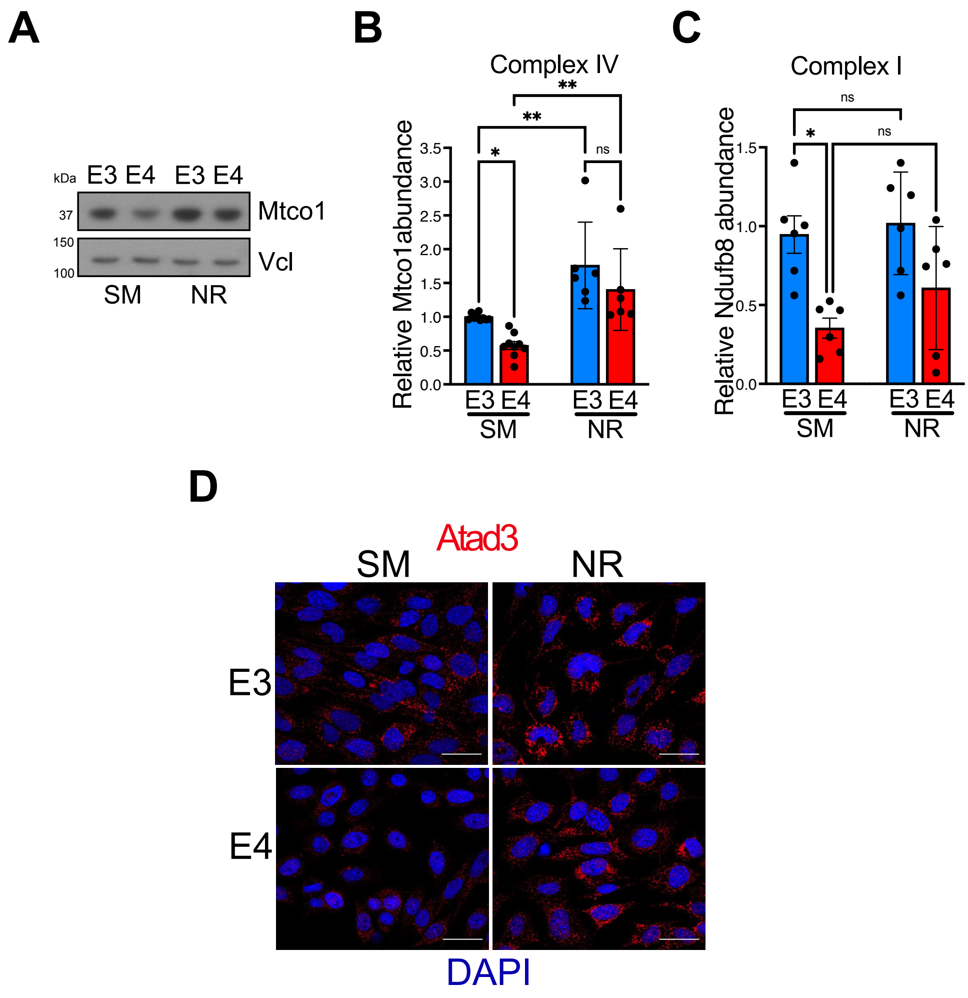


**Figure S7. A nutrient restricted medium increases respiratory chain complex IV and Atad3 in APOE4 astrocytes. A)** Representatives immunoblot of complex IV subunit MtcoI, with vinculin (Vcl) as a loading control, **B)** Quantification of Mtco1 in APOE3 and APOE4 astrocytes from n = 6 experiments, Mtco1 abundance in E4 cells is higher in NR than SM, p = 0.003. **C)** abundance of respiratory complex I subunit Ndufb8 and complex I in-gel activity in APOE4 astrocytes after 24 hours exposure to a nutrient restricted medium (NR) with 0.6 mM β-hydroxybutyrate, versus standard medium (SM). Several of the cell protein samples are the same as those that revealed an increase in Opa1L with nutrient restriction (Figure 7E). **D)** APOE3 and APOE4 astrocytes were cultured on standard medium (SM) with 25 mM glucose and 1 mM pyruvate or on a cholesterol lowering nutrient restricted regime, NR (i.e. without glucose or pyruvate and with 0.6 mM β-hydroxybutyrate). Cells were immunostained for Atad3, and the nuclei labeled with DAPI. Scale bar = 50 μm.

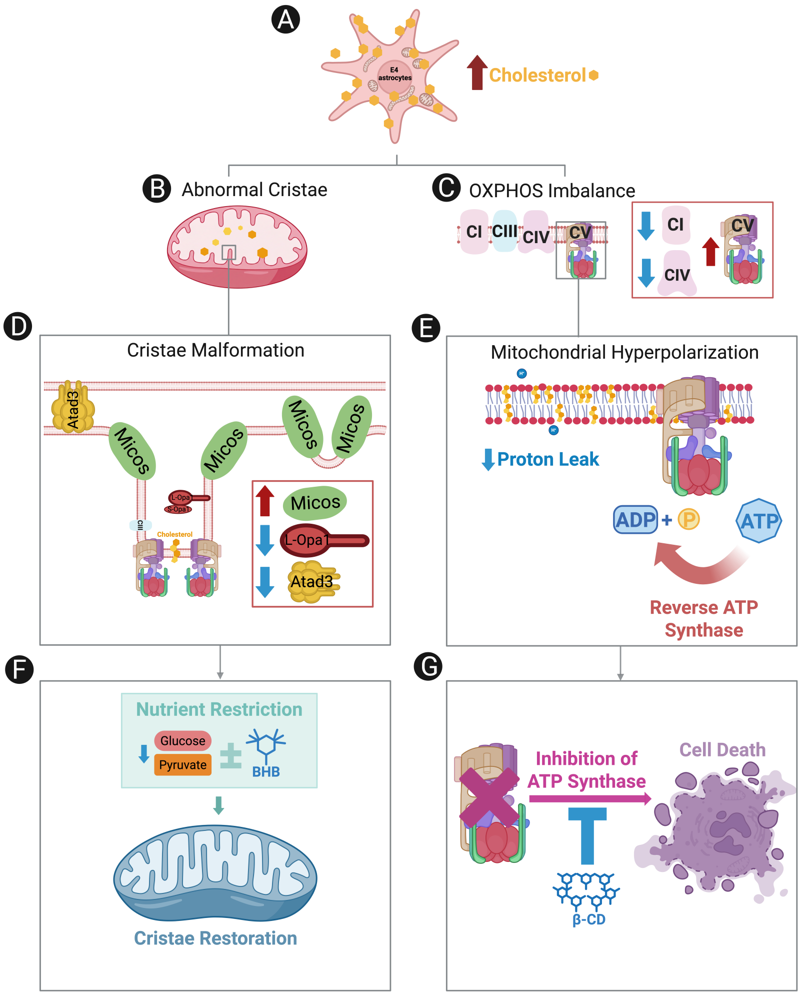


**Figure S8. Elevated cholesterol in *APOE4* astrocytes drives mitochondrial structural and functional abnormalities.** **A**) APOE4 astrocytes accumulate excessive intracellular cholesterol. **B**) Elevated cholesterol disrupts mitochondrial cristae architecture leading to **C**) impaired OXPHOS. **D**) Cristae disorganization is associated with altered levels of the cristae junction complex Micos, L-Opa1, and Atad3. **E**) The altered inner mitochondrial membrane results in reduced proton leak and increased reverse ATP synthase activity. **F**) Nutrient restriction, via glucose and pyruvate withdrawal, with or without β-hydroxybutyrate supplementation, restores normal cristae morphology. **G**) Inhibition of ATP synthase with oligomycin that selectively induces cell death in APOE4 astrocytes is rescued by methyl β-cyclodextrin treatment (β-CD).

| Antigens | Manufacturer | Catalogue number | Dilution (WB) | Dilution (ICC) |
|---|---|---|---|---|
| | | **Primary antibodies** | | |
| ABCA1 | Novus | NB400-105SS | $1000^{-1}$ | |
| APOE | Santa Cruz | sc-390925 | $1000^{-1}$ | |
| APOE4 | Novus | NBP1-49529SS | $1000^{-1}$ | |
| ATAD3 | Proteintech | 16610-1-A | $1000^{-1}$ | $200^{-1}$ |
| ATP5B | Proteintech | 17247-1-AP | $5000^{-1}$ | |
| HMGCR | Santa Cruz | sc-271595 | $1000^{-1}$ | |
| Mitofilin | Proteintech | 10179-1-AP | $1000^{-1}$ | $200^{-1}$ |
| MTCO1 | Abcam | ab14705 | $5000^{-1}$ | |
| NDUFB8 | Abcam | ab110242 | $1000^{-1}$ | |
| OPA1 | BD | 612607 | $1000^{-1}$ | |
| PHB1 | Cell Signaling | 2426 | $1000^{-1}$ | |
| PHB2 | Cell Signaling | 14085 | $1000^{-1}$ | |
| TOMM20 | Proteintech | 11802-1-AP | $5000^{-1}$ | |
| UQCRC2 | Abcam | ab14725 | $1000^{-1}$ | |
| VCL | Santa Cruz | sc-73614 | $10000^{-1}$ | |
| | | **Secondary antibodies** | | |
| Goat Anti-Mouse IgG (H+L), HRP | Promega | W4021 | $5000^{-1}$ | |
| Goat Anti-Rabbit IgG (H+L), HRP | Promega | W4011 | $5000^{-1}$ | |
| Alexa FluorTM 488 goat anti-mouse IgG (H+L) | Invitrogen | A11029 | | $450^{-1}$ |
| Alexa FluorTM 555 goat anti-rabbit IgG (H+L) | Invitrogen | A13572 | | $450^{-1}$ |

**Table S1. List of antibodies used in this study.** The table summarizes all antibodies employed for immunodetection. The Primary Antibody column indicates the antibody used to detect the target antigen. The Secondary Antibody column specifies the antibody used for detection of the primary antibody, including conjugates (e.g., HRP, Alexa Fluor). Additional details such as manufacturer, catalogue number and dilution are provided for reproducibility.
